# Beach environmental DNA fills gaps in photographic biomonitoring to track spatiotemporal community turnover across 82 phyla

**DOI:** 10.1101/680272

**Authors:** Rachel S. Meyer, Teia M. Schweizer, Wai-Yin Kwan, Emily Curd, Adam Wall, Dean Pentcheff, Regina Wetzer, Eric Beraut, Alison Young, Rebecca Johnson, Robert K. Wayne

**Author notes:** Co-first author. Colorado State Univ Dept of Biology, Fort Collins CO 80523 RS Meyer and TM Schweizer should be considered joint first author. Corresponding author: 001-206-351-7997. Author Contributions: RSM, TMS, EC, AY, RJ, and RKW designed the study, with WK and AW designing and performing additional analyses. RSM, TMS, EC, and RKW generated the eDNA data, DP and RW contributed DNA reference data and analyzed taxonomic results. RSM, EB and WK managed the data. All authors contributed to analyses and to writing the manuscript.

## Abstract

Environmental DNA (eDNA) metabarcoding is emerging as a biomonitoring tool available to the citizen science community that promises to augment or replace photographic observation. However, eDNA results and photographic observations have rarely been compared to document their individual or combined power. Here, we use eDNA multilocus metabarcoding, a method deployed by the CALeDNA Program, to inventory and evaluate biodiversity variation along the Pillar Point headland near Half Moon Bay, California. We describe variation in presence of 13,000 taxa spanning 82 phyla, analyze spatiotemporal patterns of beta diversity, and identify metacommunities. Inventory and measures of turnover across space and time from eDNA analysis are compared to the same measures from Global Biodiversity Information Facility (GBIF) data, which contain information largely contributed by iNaturalist photographic observations. We find eDNA depicts local signals with high seasonal turnover, especially in prokaryotes. We find a diverse community dense with pathogens and parasites in the embayment, and a State Marine Conservation Area (SMCA) with lower species richness than the rest of the beach peninsula, but with beta diversity signals showing resemblance to adjacent unprotected tidepools. The SMCA differs in observation density, with higher density of protozoans, and animals in Ascidiacea, Echinoidea, and Polycladida. Local contributions to beta diversity are elevated in a section of East-facing beach. GBIF observations are mostly from outside the SMCA, limiting some spatial comparisons. However, our findings suggest eDNA samples can link the SMCA sites to sites with better GBIF inventory, which may be useful for imputing species from one site given observations from another. Results additionally support >3800 largely novel biological interactions. This research, and accompanying interactive website support eDNA as a gap-filling tool to measure biodiversity that is available to community and citizen scientists.

## 1. Introduction

The 6th mass extinction now underway is projected to affect a million species (Diaz et al., 2019), and consequently, scientists and the public need to catalyze the development of next-generation biodiversity monitoring strategies for a comprehensive inventory of biodiversity (National Research Council, 2001). Our interest is in a strategy that activates a public constituency with access to the latest digital and molecular tools and an understanding of the data and results so they can innovate solutions together with the scientific community. Volunteers for citizen and community science outnumber professional scientists 18 to 1 (Groom et al., 2017), and have contributed a substantial amount of observational data points that are now used to track biodiversity variation over space and time (Bird et al., 2014). One major source of observations is the iNaturalist community science platform [Cal Academy of Sciences (CAS) and National Geographic Society], which has exceeded 12 million ‘research grade’, or photographed and validated, records. For certain species and localities, these observations comprise the bulk of data points over other occurrence data, such as museum and voucher specimens. iNaturalist and other volunteer-collected species occurrence records (such as eBird) help populate the Global Biodiversity Information Facility (GBIF; Robertson et al., 2014), which also includes collection and observation data curated by thousands of institutions (n=1332 ‘publishers’ as of January 11, 2019). Citizen and community scientists are eager to be data contributors and deepen their involvement in knowledge assessment and possible conservation actions, (Hecker et al., 2018). Everyone is impacted by the challenges of biodiversity loss.

Environmental DNA (eDNA) presents an attractive method for obtaining alternative biodiversity data. The method is considered non-invasive as small amounts of soil, sediment, or water are sufficient for DNA extraction, which can be easily collected by volunteers or passively collected with unmanned devices. eDNA can be subjected to multilocus metabarcoding and next-generation sequencing, producing snapshots that span all kingdoms of life. However, because we don’t have sufficient DNA barcodes for all species, sometimes there are insufficient data to resolve taxonomy or avoid mismatches. DNA research in the environment is also limited by the non-even distribution of DNA from taxa (especially inshore and terrestrial environments; O’Donnell et al. 2017), where small bodied and more evenly distributed species have better detectability over large mobile organisms (Zinger et al. 2019a). Therefore, eDNA-based biodiversity inventories may benefit from complementation with other evidence-based observations such as photos or specimens. Nonetheless, the sheer data-richness of eDNA may elucidate robust metacommunities with dynamics that inform ecosystem management (e.g. Peters et al., 2019) and that may even allow imputation about the presence of larger organisms from their associated smaller taxa comprising their holobiomes (i.e. the total genomes in and on a eukaryotic organism, possibly functioning together as evolutionary units; Guerrero et al., 2013). However, as a nascent field, this topic has hardly been explored.

Photographic observations present challenges that systematic collection of eDNA may help resolve. Diversity estimates are limited to where people choose to go, to what people decide to photograph or sample, or to where camera traps are installed. Further, by the nature of photographs, it is mainly relatively large or morphologically apparent, diurnal organisms. How can eDNA surveys augment or predict taxa in observational databases? If eDNA provides a next-generation biodiversity assessment tool, how can it be integrated into biomonitoring by the public and scientists? By using eDNA samples to fill information gaps and stimulate broader human observation, can we mitigate taxon (e.g. plant blindness; Allen, 2003) and visitation bias?

This study presents an analysis of eDNA metabarcoding collected through community science effort of the CALeDNA program (www.ucedna.com; Meyer et al., 2019). Our analysis focuses on an inventory of biotic communities on the Pillar Point headland California beach that has been previously been intensively surveyed by the California Academy of Sciences (CAS) Citizen Science team (https://www.inaturalist.org/projects/intertidl-biodiversity-survey-at-pillar-point). The beach has a marshland and harbor graded C-F for water quality on the Heal the Bay Beach Report Card (2016; healthebay.org), signaling it is polluted and potentially under threat. It also has a State Marine Conservation Area (SMCA) with restricted fishing, that is difficult to access by land, which received grades A-C. It remains unclear how biologically different these areas are. We compare eDNA results to human observation records (GBIF; largely including iNaturalist research grade observations) to determine their complementarity and the value of each to fill gaps where data are limited across space and time. We find eDNA and photographs have little overlap, with eDNA covering lower trophic. Systematic collection of eDNA can describe differences spatial and temporal biodiversity variation of the beach, and provides a rich view of the community ecological network, including holobiomes. Even though this is only a single case study, our results suggest that eDNA may be useful to find surrogate sites where GBIF data is readily collected, which may be useful proxies for diversity in sites that are difficult for the volunteer community to monitor with photographs. With community empowerment as the goal, results in this paper are paired with interactive web pages for the public, who are potential eDNA adopters, to explore (https://data.ucedna.com/research_projects/pillar-point<u>?</u>).

## 2. Materials and Methods

Detailed methods including commands for running programs are in the Supplemental Methods.

### 2.1. Sample collection for eDNA

Samples of soil, sand and sediment for eDNA analysis were collected at Pillar Point by a total of fifteen community scientists and three UCLA researchers during low-tide windows of three days: February 8, 2017, April 29, 2017 and April 30, 2017. Our definition of community scientist is a volunteer participating in a science project, which could be designed and empowered by non-professional scientists or organized by professional scientists [see Meyer and Drill (2019) for further discussion]. The February sampling date coincided with the monthly Pillar Point survey organized by CAS Citizen Science Academy staff, which selects seasonally low tides for observation. In February, most eDNA samples were only collected near or above the mean high water line (aka mean high tide mark; MHT) because of permit restrictions. Most sites below the MHT line were sampled in April, as were all SMCA samples. Collections were made under permit MBNMS-2017-019 issued by the NOAA Office of National Marine Sanctuaries. Sample metadata are provided in Table S1.

For the eDNA collection, sample collectors were organized into groups to cover different areas of the Pillar Point headland. Sample collectors then spread out and selected sites that looked like a typical representation of the local environment. Surface soil, sand, or submerged sediment were collected following the CALeDNA community science program instructions, which required fresh gloves to be worn at each sampling site, three ‘biological replicates’ of 2 mL cryotubes to be filled with substrate approximately 30 cm apart from each other. Sampling was limited to the exposed (top 3 cm layer) surface or submerged sediment no deeper than could be reached at arm’s length. Collectors used a Kobo Toolbox phone webform (https://ee.kobotoolbox.org/x/#Y1sl) to record sampling time, geolocation, photographs of the sampling site, and other metadata. Metadata for each sample are available through data.ucedna.com. Photographs were later used to confirm environmental metadata (Table S1) for the sites. All collection instructions, data and photographs are published online in the CALeDNA database at www.ucedna.com.

### 2.2. Assembling observation records

GBIF records from all contributors were downloaded September 4, 2018 from GBIF using drawn polygons on a map of Pillar Point (coordinates and download DOI in Appendix 1). eBird observations were removed because they were concentrated only in the embankment. A total of 13,924 occurrence records were retained. The polygons correspond to the headland embayment, unprotected reef area, and SMCA protected area (data in Table S2).

### 2.3. eDNA metabarcoding library preparation and sequencing

Subsamples of equal mass of all three biological replicates were pooled and homogenized. A ∼0.25g amount of the subsample was extracted with the Qiagen DNeasy PowerSoil Kit according to manufacturer’s instructions (Qiagen, Valencia, CA, USA). Kit details provided by the manufacturer demonstrate retrieval of amplifiable DNA from bacteria, fungi, algae, and animals. A blank PowerBead tube was extracted alongside each batch of ∼10 samples as a negative extraction control. Eight DNA extraction controls were pooled into a single sample used in all PCR amplifications.

Sample DNA was amplified for targeted taxa using four primer sets based on the following universal primers with a Nextera adaptor modification: MiFish 12S (Universal Teleost; Miya et al. 2015), 16S (515D and 806R; Caporaso et al., 2012), 18S (V8-V9; Amaral-Zettler et al., 2009; Bradley et al., 2016), and the partial sequence of Cytochrome Oxidase Subunit 1 (CO1; mlCOIintF and jgHCO2198; Leray et al., 2013). Primer sequences are included in Supplemental Methods. Three PCR replicates were run per amplicon per sample, and these were pooled and converted to Nextera libraries (details in Supplemental Methods), and sequenced using a MiSeq Illumina Next Generation Sequencer with Reagent Kit V3 (2 x 300 bp) at a goal depth of 25,000 reads in each direction per marker per sample. The original run produced reverse reads with lower than standard quality, and so the run was repeated. Both runs were analyzed separately and resulting tables were merged.

### 2.4. eDNA classification

Demultiplexed library Fastq files were processed with the Anacapa Toolkit (archived version (doi:10.5281/zenodo.3064152)(github.com/limey-bean/Anacapa; Curd et al., 2019) using default parameters. In brief, reads were processed first by cutadapt (Martin, 2011) to remove adapters and any 3’ primer reverse complement. Reads were quality trimmed with the FastX-Toolkit (http://hannonlab.cshl.edu/fastx_toolkit/) and only reads with a minimum length over 100 bp were retained. Cutadapt was used to sort reads by primer and then used to remove primers. Read bins were denoised, dereplicated and merged or retained as paired unmerged, forward-only or reverse-only reads using dada2 (Callahan et al., 2016). Chimeras were additionally removed using dada2. This produced amplicon sequence variants (ASVs) that were then processed using Bowtie 2 (Langmead and Salzberg, 2012) to query custom reference databases (described in the following paragraph) and determine up to 100 reference matches with a minimum percent coverage of the sample read set to 70% for CO1, 80% for all others and a minimum percent identity of 70% for CO1, and 90% for all others. Details on how these settings were chosen are in Supplemental Methods and Table S3. Reads per taxon were compiled for each marker using a minimum bootstrap classifier confidence (BCC) = 60 for 16S, 12S, and 18S, and a minimum BCC = 70 for CO1.

Reference databases for 16S, 18S, CO1, and 12S were generated with the CRUX step of the Anacapa Toolkit (Curd et al., 2019; DOI: https://doi.org/10.5061/dryad.mf0126f) that queried NCBI nr/nt databases. The CO1 database was subsequently modified by appending all BOLD CO1-5P sequences, accessed via the BOLD API on September 24, 2018 and comprising 5.08 million sequences, and also appending 847 recently generated invertebrate CO1-5P barcode sequences from the LACM DISCO program (subsequently published in BOLD with the same identifiers). The CO1-5P region is the most commonly used CO1 locus, described first by Folmer et al (1994) and it includes our amplified region. These reference databases are permanently archived [in DRYAD – link pending].

BOLD and NCBI use different classifications systems for higher taxonomy (order to superkingdom). Results tables originally had taxon strings that could be a mixture of classifications. We united them using the higher classification by Ruggiero et al. (2015; online version) for all results tables and merged rows with identical names by summing column read counts. This procedure was only used for decontaminated results.

For 16S, 18S, and 12S markers, we statistically removed reads and taxa considered contaminants using the program Decontam version 1.1.2 (Davis et al., 2018) with prevalence method and the threshold setting of 0.1. Taxa with only a single total read were subsequently removed in the final results (Tables S4-S7).

For CO1, because there were few taxa that overlapped between real samples and extraction or PCR negative controls, results tables were decontaminated simply by removing any taxon that was present in one or more reads in controls. The extraction and PCR negative controls contained a total of three taxa, two being single reads in either control or real sample, and one that did occur as 9 reads in a control and in more reads in four real samples, which was Nutricola tantilla Gould, a very small clam. We did not include this taxon in results, and note it was not in GBIF Pillar Point data but is known from California (GBIF.org). Singletons were also subsequently removed.

Using the CO1 results, we further examined possible contaminants by heeding Zinger et al., (2019b) who describe the various contamination issues, specifically, the risk of index hopping among samples in the same MiSeq run. We looked for cross-contamination between the Pillar Point samples, which were the ∼54% of the identified reads on the MiSeq run, and California Vernal Pool samples which were largely the remainder of the run, and found only 43 taxa overlapped, 15 of which were singletons in the Pillar Point samples, and 34 of which were soil-dwelling Ascomycota or amoeba that could theoretically occur in both habitats (Table S6). Not one read of known vernal pool-specific animals, which each had >2500 reads in the vernal pool samples, were recovered in Pillar Point samples (e.g., Gammarus cf. fossarum, or Alona, which are usually found in ephemeral freshwaters and known from vernal pools; Keeley and Zedler, 1998; Hupalo et al., 2018). We also counted the amount of PhiX in sample libraries using BBDuk (BBMap – Bushnell B. – sourceforge.net/projects/bbmap/), and found it was (<8e-05) in each library. We found these low occurrences of contamination from extraction or index-hopping suggested further decontamination steps were unnecessary. The Anacapa output results including the non-Pillar Point samples, and files detailing sequences themselves and other details of taxonomy assignment confidence, are provided for CO1 in Table S6.

Alpha diversity analyses used a stricter filter of a minimum of five reads to reduce some inflation effects from ASVs. All analyses requiring rarefaction used settings retaining 2000 reads for 18S or CO1, and 4000 reads for 16S, aiming for a cutoff above the linear growth zone of the curve. Rarefaction curves used to inform the sequence number for rarefaction are shown in Figure S1.

### 2.5. Global biotic interactions (GloBI)

We downloaded all of the open access GloBI species interactions database (Poelen et al., 2014; http://globalbioticinteractions.org) on September 2, 2018, totaling 3,293,470 records. These data were used in two tests. First, we retrieved all the species that had 10 or more observations from the Intertidal Biodiversity Survey at Pillar Point in iNaturalist (https://www.inaturalist.org/projects/intertidal-biodiversity-survey-at-pillar-point?tab=species), and searched for all GloBI records of interactions involving these frequently observed species (<u>Biotic Interactions</u>). We screened the list of interacting taxa for overlap with GBIF species and eDNA species and reported these. Second, we summarized all GloBI interactions to family level, and recorded the interaction type. We used the major categories of “eats” or “interacts_with” to tally the frequency with which the families detected in eDNA or GBIF are on source or sink for these interactions, and used these sums in Chi-Square tests in R (version 3.5.0) to test for overrepresentation of interaction source or sinks in eDNA and GBIF results.

### 2.6. Statistical analyses for diversity and enrichment

The 16S, 18S, and CO1 results (Tables S4-S7) were used to generate separate alpha diversity and beta diversity plots, and used to calculate statistics including Local Contributions to Beta Diversity (LCBD) and Analysis of Similarity (ANOSIM). To prepare results for these tests and plots, decontaminated Anacapa results tables and metadata were converted to Phyloseq objects (McMurdie and Holmes, 2013) using the convert_anacapa_to_phyloseq function in the Ranacapa version 0.1.0 (Kandlikar et al., 2018) R package in R version 3.5.0. Phyloseq and microbiomeSeq R package version 0.1 (https://github.com/umerijaz/microbiomeSeq.git) were used to calculate richness, Simpson, and Shannon alpha diversity and test for significant differences among groups with ANOVA set to P=0.05. Plots were made using ggplot2 (Wickham 2009). Full scripts are provided in Supplemental Methods.

Beta diversity as relative abundance barplots and LCBD statistics were generated in R using the addition of the adespatial (multivariate multiscale spatial analysis) package version 0.3.2 function selecting the Hellinger dissimilarity coefficients method (Dray et al., 2016). ANOSIM beta diversity statistics and distance plots were computed using both Jaccard and Raup-Crick dissimilarity indices (Raup and Crick, 1979). Both treat the data as presence/absence but Raup-Crick considers underlying alpha diversity, as calculated in the Vegan vegdist method (Chase et al., 2011; Oksanen et al. 2018) in Phyloseq. Principal Coordinate Ordinations were made using the Jaccard method.

We performed density enrichment tests in R using the following approach. Taxon results tables were summarized by class and converted to sample presence/absence tables. Kruskal-Wallis rank sum tests (Hollander, et al., 2013) were performed to identify classes with significant differences among zone groups. Then, significant classes were subjected to the post-hoc Dunn test (Dunn, 1964) using the Benjamini-Hochberg (Benjamini and Hochberg, 1995) correction for multiple testing.

### 2.7. Community ecological network analyses

Network analysis was performed using the SpiecEasi version 1.0.2 package (Kurtz et al., 2015; Tipton et al., 2018) in R with the 18S, 16S, and CO1 results tables summarized to the highest resolution classification of family. We did not include 12S because the taxon list was short. Family-level results tables were filtered to retain only taxa minimally present in 15% of the sites. Results from each marker were processed separately, and then 16S was co-analyzed with each of the dominantly eukaryotic markers 18S and CO1. Settings and commands are in Supplemental Methods, as are commands for plotting networks using Phyloseq and iGraph (Csardi and Nepusz, 2006). Networks were additionally plotted as interactive figures using Flourish Studio (https://app.flourish.studio; London, UK), given a stable DOI, and were hosted on our Pillar Point project website (https://data.ucedna.com/research_projects/pillar-point?). Interactions were also deposited in GloBI (https://github.com/beraute/Pillar_Point_16S_18S, https://github.com/beraute/Pillar_Point_CO1_16S). We asked what proportion of observed eDNA network interactions were previously published, and tested this by comparing 18S network taxa with 10 or more edges (highly networked) to interactions published in the GloBI database (Section 2.5). Lists were assembled and compared in Microsoft Excel.

### 2.8. Reduction of holobiome effects

For the 18S results, we tested the influence of DNA swamping from organisms and their associated community (holobiome) on local contribution to beta diversity (LCBD) scores. We screened the 18S rarified DNA results for taxa with >50% proportion of the read abundance. Community ecological networks (Section 2.7) were mined for these organisms and their linked taxa, and all these taxa were filtered out to produce new results tables. These holobiome-filtered tables were used in LCBD calculations as described in Section 2.6.

### 2.9. GBIF and eDNA comparisons

GBIF and eDNA results use different classification systems. The classification at each hierarchical level was converted to a common NCBI-style taxonomy using the Global Names tool (http://globalnames.org). This tool has been shown to increase the success of cross-mapping taxon names to up to 90% across diverse databases (Patterson et al., 2016); in accordance with this number, under 10% of taxa had to be dropped because they could not be converted. This slightly reduced set of converted names is presented on the web platform.

### 2.10. Development of the web interface and comparative summary statistics

A web platform was created linking Squarespace, Amazon Web Services, and Heroku. Numerous data visualization tools such as leaflet were used to create the user interface, and images were scraped from Encyclopedia of Life (eol.org), GBIF (gbif.org) iNaturalist and Wikipedia. All scripts to generate the pages are open source and available on Github at https://github.com/UCcongenomics/caledna.

## Results

### 3.0. Orientation

Pillar Point headland contains a 1 km East-facing stretch that spans a small protected marsh with agricultural runoff feeding into an embayment that contains a harbor with primarily fishing and recreational crafts (Figure 1). The marsh supports a diversity of seabirds and shorebirds. The entire area is bounded by a stone embankment (breakwater). A 0.4 km stretch of South-facing ‘outer’ beach has an extended 0.3 km tidepool accessible during very low tides that receives heavy recreational traffic and visits from school groups as well as commercial and recreational collection of mollusks and fish. North of the tidepools on the West ‘outer’ beach (Figure 1) is a 0.8 km stretch that includes the Pillar Point State Marine Conservation Area (SMCA). The SMCA places limits on recreational and commercial fishing, but not access for recreational users (Marine Life Protection Act of 1999).

**Figure 1.**
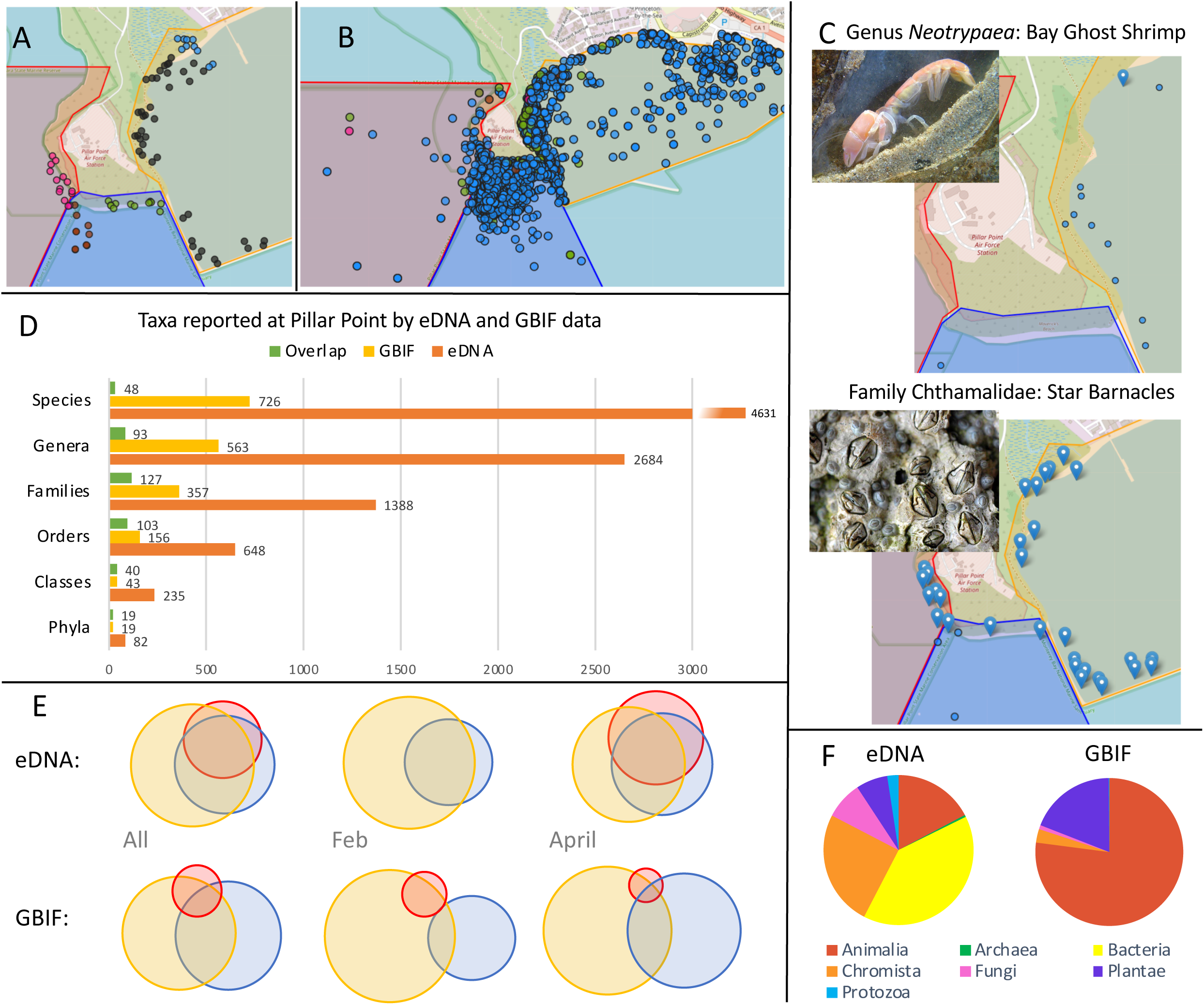
A-C. Map of the Pillar Point beach with highlighted polygons designating embayment (orange), unprotected (blue), and protected (red) areas used to group sites. A. The 88 eDNA sites are color coded by zones. From top right, marsh runoff (blue), unprotected inner beach (grey), unprotected outer beach (green), unprotected tidepools (red), and protected outer beach (SMCA; pink). B. The 13,942 GBIF observations, color coded by kingdom, with Animalia (blue), Plantae (green), Fungi (pink), and Chromista (red). C. Examples of website functions for taxa contained in both eDNA and GBIF datasets. eDNA sites where a taxon was found are drop pins and GBIF sites where a taxon was found are dots. Bay ghost shrimp are commonly photographed but eDNA only detected them once. Star barnacles were only rarely documented in the unprotected tidepools with photographs, but they are found all over the area by eDNA. D. The total number of unique taxonomic lineages in eDNA, GBIF, or in common. The common taxa are also included in the plotted totals of each separate database. E. Venn diagrams of overlapping and unique taxa found in eDNA and GBIF datasets for the total data (All), for the February 2017 collection for eDNA and the month of all February GBIF observations made over all years for which data was available, and for the April 2017 eDNA collection and April GBIF observations made over all years data was available. Colors for polygons are consistent with A and scaled to the relative number of taxa out of the total taxa included in that particular diagram. F. Pie charts of the proportion of unique taxa inventoried by each method. Photo credits: Monterey Bay Aquarium and Wild Kratts Wiki.

We examined the inventory of taxonomic biodiversity at Pillar Point beach from eDNA results and GBIF records, grouping data in several ways. The beach areas were divided into three polygons drawn on a map: 1) unprotected “embayment”; 2) “unprotected” exposed beach and tidal pools; and 3) “protected” exposed beach of the SMCA (Figure 1A, 1B). We also divided the beach into five zones: 1) area exposed to marsh runoff; 2) unprotected inner beach; 3) unprotected outer beach; 4) unprotected tidepools; and 5) protected outer beach (SMCA). The total taxa observed (Figure 1D) were used to compare biodiversity among the polygons. We added metadata for each sampling site that included the month collected, the position relative to the mean-high-tide (MHT) line, and the substrate. Comparative analyses of eDNA (Tables S4-S7) and GBIF results (Appendix 1; Table S2) assessed species in common, differences in estimates of taxon distribution, richness, beta-diversity within and among areas of the beach (Figure 1C-F). We also evaluated the kinds of biotic interactions between taxa in both types of inventory. As a community science project, we presented interactive results to engage non-scientists and scientists alike. Those resources are available in this web platform: https://data.ucedna.com/research_projects/pillar-point and are permanently archived in Dryad and Zenodo [PENDING MANUSCRIPT ACCEPTANCE]. In the Results, we highlight the website features intended for self-driven evaluation of eDNA and GBIF biomonitoring. The web platform explorable features are described in Table 1.

**Table 1.**
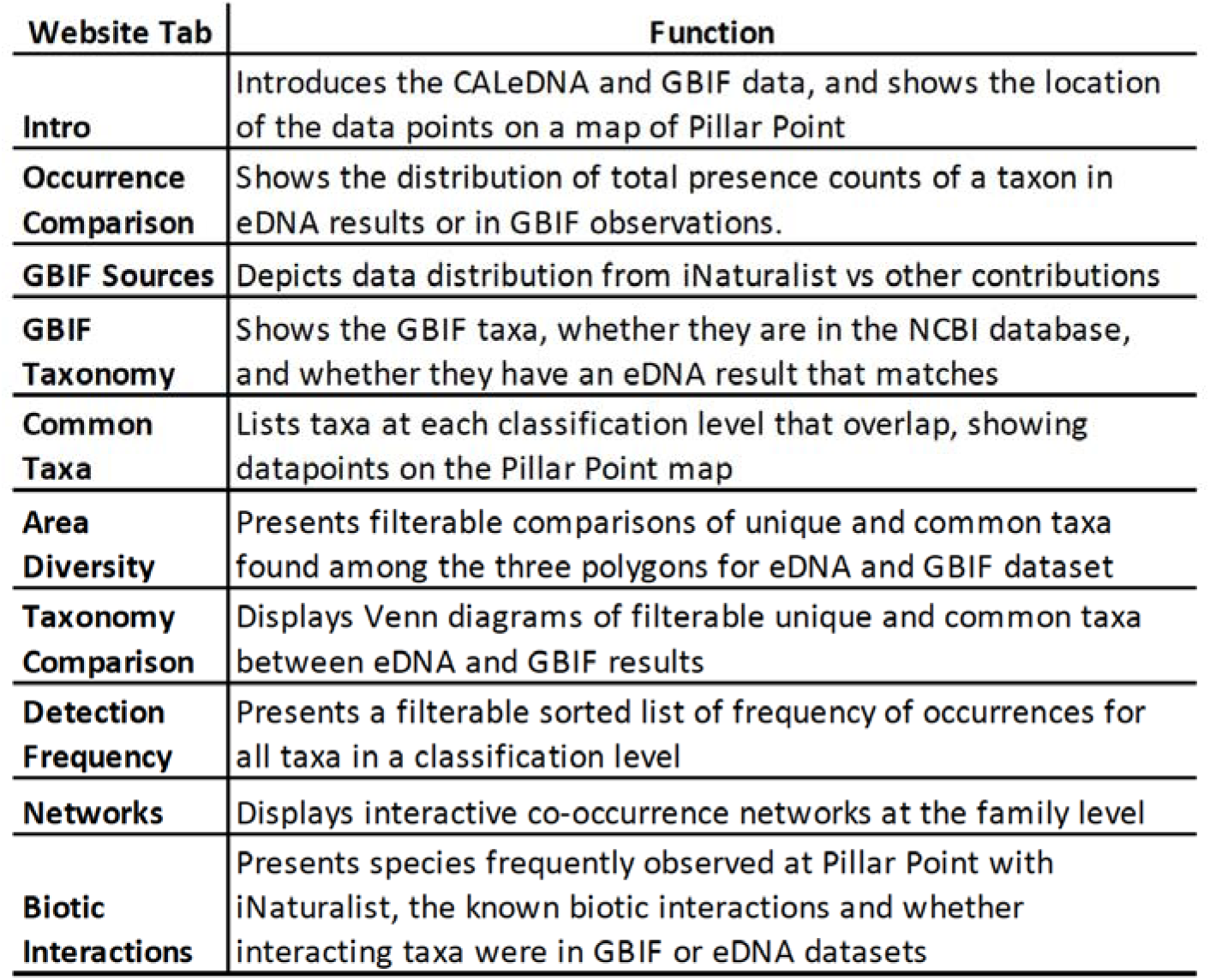
Features of the project website, https://data.ucedna.com/research_projects/pillar-point

### 3.1. eDNA taxonomic inventory and richness

Multilocus metabarcoding for four gene loci targeted Bacteria and Archaea (16S), animals and algae (CO1), eukaryotes (18S), and fish (12S). CO1 and 18S partially overlap with regard to organisms sequenced. Libraries for 88 sites and negative controls were sequenced in two runs: 3.34 Gb of fastq data were generated in the first run (170818_300PE_MS1) and 12.25 Gb were generated in the second run (171006_300PE_MS1)(NCBI SRA PENDING ACCEPTANCE). Results were output from the Anacapa Toolkit (Curd et al., 2019) summarized to the Least Common Ancestor, and then taxa were removed that were found in negative controls as well as taxa with only one read across all samples, yielding 1468 total taxa for CO1, 2689 for 18S, 2593 for 16S, and 43 for 12S (Tables S4-S7). The total unique taxa assigned to kingdoms across all eDNA results were 1132 Animalia, 27 Archaea, 2,533 Bacteria, 1,588 Chromista, 516 Fungi, 433 Plantae, and 154 Protozoa, with 3 unassigned to a kingdom.

Alpha diversity, defined by Whittaker (1972), is a measure of species richness of a place. DNA metabarcoding data has been routinely used to estimate alpha diversity for a decade (Fonseca et al., 2010), and can be used for measuring genetic diversity as well as taxonomic diversity. We focus our biodiversity analyses on taxonomic diversity, presenting alpha diversity analyses and its interpretation in Appendix 2, but include an assessment of how genetic and taxonomic diversity are related (Table S8). In taxonomic alpha diversity analyses, we found seasonal and spatial differences in both prokaryotes and eukaryotes (Appendix 2; Figure S2).

We asked how much sampling month (February or April) changed the families detected. Across the total 1,388 families detected by the four markers, the average frequency a family was found was 18% (16 out of the 88 sites). The Pearson correlation (ρ) between the normalized frequencies of these 1388 families observed in February with April was 0.81, but in repeating the test using a simple tally of whether families were observed at all in February or April, the correlation was ρ=0.27. These results, taken together with the significant spatial and temporal differences found in alpha diversity analyses (Appendix 2) support eDNA signals are spatially local and temporally restricted.

### 3.2. eDNA-based beta diversity

Analysis of Similarities (ANOSIM) beta diversity tests were performed to evaluate differences in the groupings of polygon, month, position in relation to the MHT line, and zone. Raup-Crick dissimilarity (Raup and Crick, 1979) detected fewer significant groupings than the Jaccard method (Table S9). Raup-Crick dissimilarity results showed that month was significant for all markers and had a large effect size: 18S (R2=0.76), 16S (R2=1), and CO1 (R2=0.53). The position relative to the MHT line was significant for 16S and 18S, but effect size was small (R2<0.08). Polygon was significant for 18S and the effect size was R2=0.35.

Jaccard dissimilarity spatial ordination analyses showed month varied predominantly along the first axis for 18S and 16S but not CO1 (Figure 2). Jaccard analysis grouped by zone showed the SMCA protected polygon, only sampled in April, clustered with other April samples from the unprotected tidepools (Figure 2). We also observed that for 16S results, inner beach samples clustered with other zones, falling either with marsh runoff samples, or as a group surrounding a dense cluster of unprotected outer beach samples. This suggests the inner beach is a zone that is heterogeneous in bacteria and archaea communities, experiencing some influence from adjacent areas as evidenced by their clustering.

**Figure 2.**
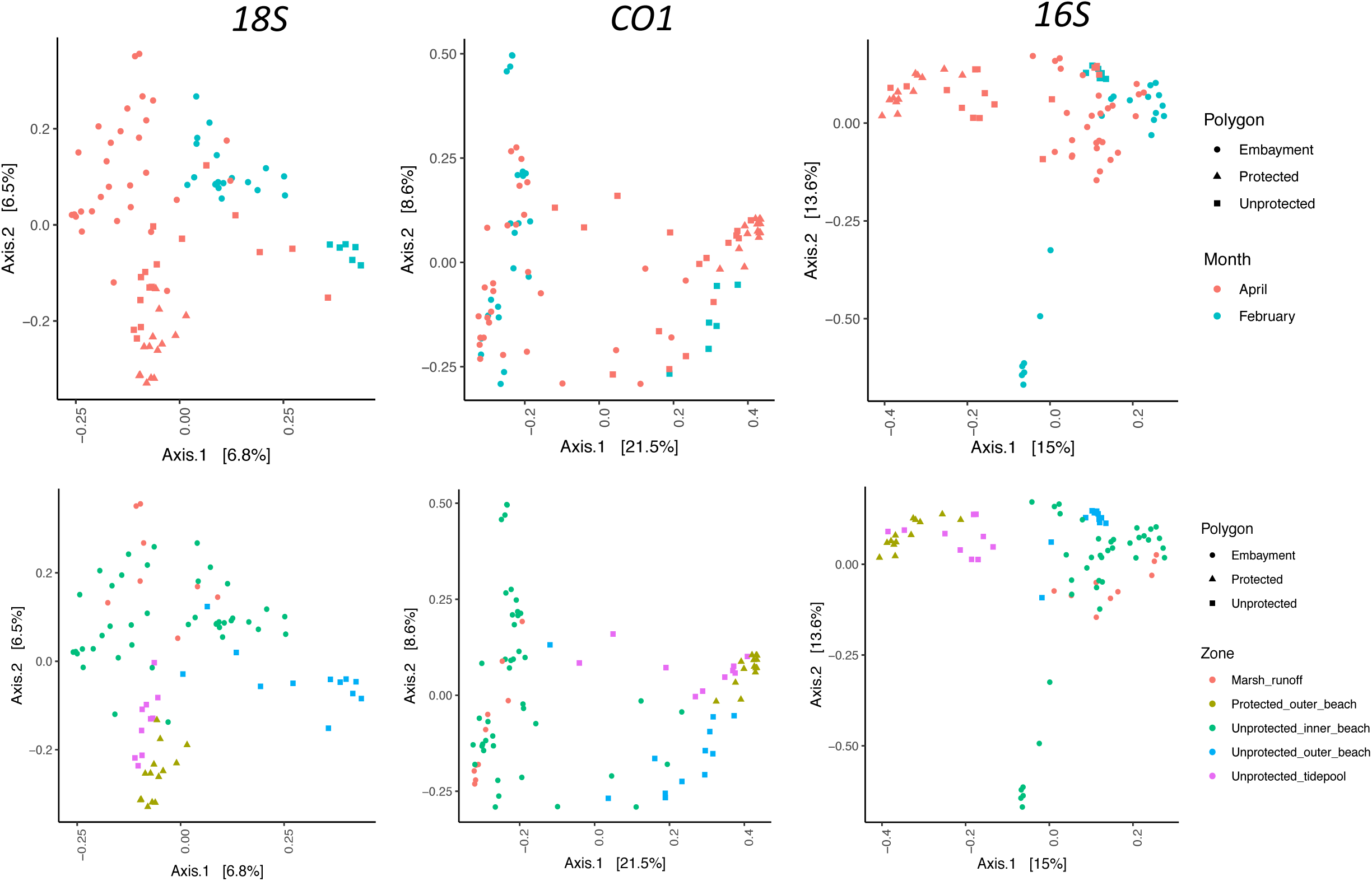
Jaccard distance principal coordinate analysis of eDNA results from the markers 16S, CO1, and 18S, color coded by zone. Results show seasonal change along Axis 1 in 18S and 16S, and similar clustering of zones across all three markers.

### 3.3. Interpreting and correcting Local Contributions to Beta Diversity

Beta diversity estimates describe the community composition and stability across localities sampled. Barplots coloring the 21 taxa with highest relative abundance show structured community composition by month, polygon, and placement relative to the MHT line (Figure 3; Figure S3).

**Figure 3.**
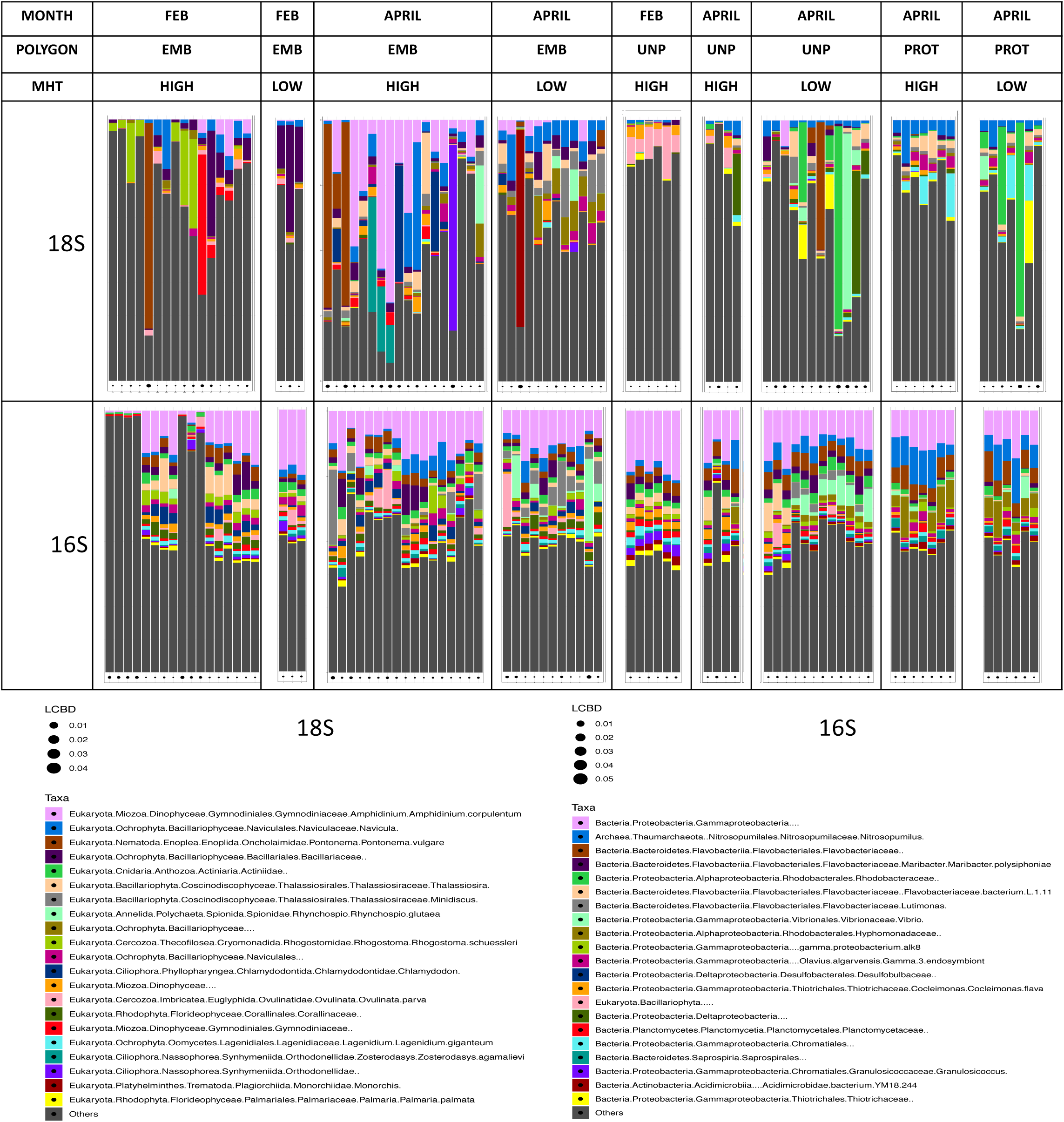
Relative abundance barplots coloring the most frequently observed 21 species. Grey bars represent the sums of all other taxa. Feb versus April indicate February and April 2017 collection dates. LCBD = Local Contributions to Beta Diversity. MHT = Mean High Tide line, Low or High indicate whether the collection was made below or above the MHT.

High relative abundance of DNA from some taxa may produce a ‘swamping effect’ that shrouds signal from other taxa (see discussion in Weber et al., 2017). In the 18S barplot, 13 samples were severely swamped with DNA (>50% of the bar; Figure 3, Figure S3), from one of several taxa, belonging to anemone, worm, ciliate, diatom, and others. Samples exhibiting DNA swamping had higher Local Contributions to Beta Diversity (LCBD) than the rest (Table S10), a measure of the uniqueness of one sample compared to the rest of its group (Legendre and Caceres, 2013). LCBD has been increasingly valued for conservation applications where researchers recommend protecting sites with extreme low and high LCBD (da Silva et al., 2018). In the 18S results of eukaryotes, there were 20 sites with significant higher or lower LCBD compared to their neighbors, 7 of which met our criteria as swamped.

We hypothesized the stability of LCBD scores was sensitive to swamping from the most abundant taxa and their associated holobiome communities, and tested this with the 18S results (Appendix 3). DNA signals from the swamping taxon were removed from results along with all of their associated taxa linked with that organism in downstream community ecological networks (Section 3.6), and then LCBD scores were recalculated. Holobiome reduction did produce lower LCBD scores for 19 samples, and only 1 of the 13 originally swamped samples remained significantly different from the rest of its group (Table S10; Figure S4). The total number of significantly different samples was reduced from 20 to 9, and all had higher LCBD than others in their group.

Eight of these nine high LCBD samples were from the cluster of the inner beach, south of the marsh. This result suggests this East-facing beach, a rarity in California, may harbor distinct communities. This inner beach is more protected from storms than outer beaches, though may be protected from marsh runoff by the breakwater. We did additional LCBD analyses to explore the similarity of values across markers (Appendix 3).

### 3.4. Enrichment tests reveal density patterns that characterize each zone

To understand biological variation at the level of spatial density, which is relevant to natural areas management, we examined which classes of taxa across kingdoms differed in their density of detection among the Pillar Point beach zones (Figure 4; Table S11). We tallied the samples within zones with DNA signal from a class, and used Kruskal-Wallis and Dunn post-hoc tests to test for enrichment in the density a class was detected. Because these results have management implications, we scrutinized our taxonomic assignments to class using phylogenetic analysis and BLAST queries to vet our trust in the correct class assignment (data not shown). We found two classes that could not be clearly delimited from other classes in their respective phyla: the class Crinoidea, which typically occurs in deeper marine environments could not be delimited from other Echinoderms. This may be due to size variation in CO1 sequences for this clade, and sequence variation shared with other organisms outside of the phylum. We also could not clearly delimit Scyphozoa from Hydrozoa, but Hydrozoa were found in all eDNA samples, and therefore have no difference in density. We therefore considered the enrichment test for putative Scyphozoa.

**Figure 4.**
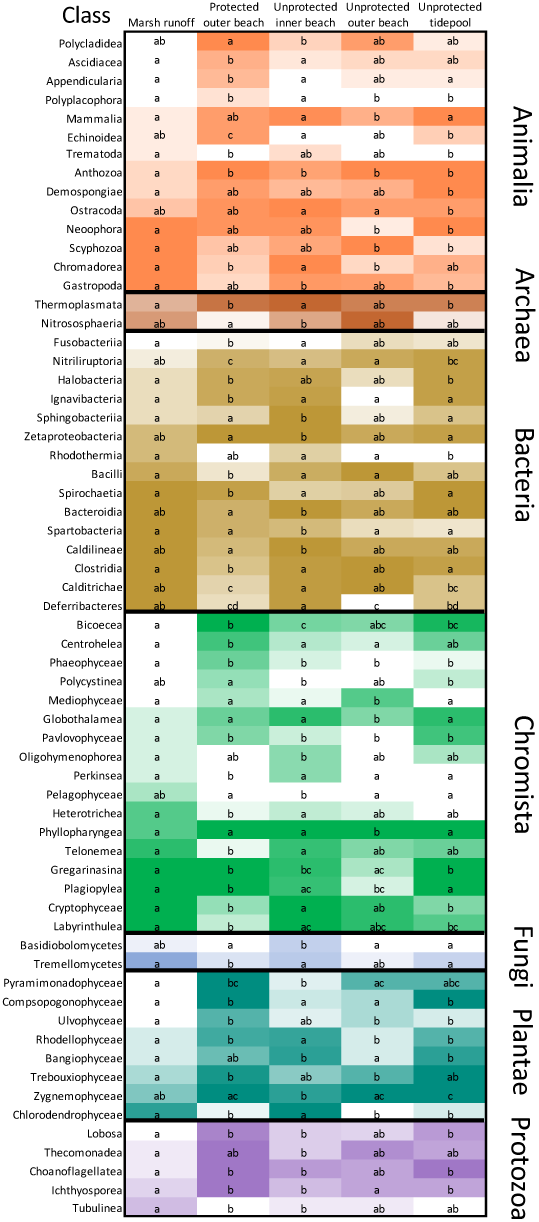
Heatmap of the eDNA-based taxonomic density within classes found in each zone, normalized to the total number of sites observed in each zone. Darker shades indicate higher density. Letters indicate significantly different groupings based on the Dunn post-hoc test, where cells with the same letter are not significantly different from each other, based on corrected p-values. Only classes with significant differences from Kruskal-Wallace test results were plotted. We note Crinoidea were removed because sequences could not be affirmatively identified to be from the class.

The SMCA was significantly more dense than all other zones in Echinoidea (sea urchins) and Ignavibacteria (a chemoheteroptrophic bacteria) and sparser in Perkinsea (parasitic Chromista causing diseases in shellfish and amphibians)(Figure 4; Table S11). The SMCA and tidepools versus other zones were significantly denser in Nitriliruptoria (often obligate halophilic bacteria), denser in Compsopogonophyceae (a red algae), sparser in Bacilli (bacteria including anthrax), sparser in Spirochaetia (ecologically diverse bacteria that includes taxa causing syphilis and Lyme disease) and sparser in Calditrichae (bacteria with few functional attributes known). The SMCA and other beaches harbored higher density of many Animalia including tunicates in Ascidiacea (sea squirts) and Appendicularia, Polyplacophora (chitons), most Protozoa and several algae (e.g. Ulvophyceae, Rhodellophyceae) compared to the marsh area. The marsh was denser in Chromadorea (roundworms), Trematoda (flukes, parasites of mollusks and vertebrates), and Gastropoda (snails and slugs). The richness of complex lifecycle parasites in the marsh suggest high presence of mobile species and genotypic diversity, such is exemplified by high trematode density where snails and migrating birds are present (Hechinger and Lafferty, 2005; Auld and Tinsley, 2015). The marsh area has rich migrating bird diversity, tracked through eBIRD (GBIF.org; see original polygon DOI download for the embayment). The unprotected inner beach, which we note had sites with elevated LCBD scores (Section 3.3), was uniquely dense in Spartobacteria, a class of both soil-dwelling and aquatic bacteria implicated in degrading algal polysaccharides (Herlemann et al., 2013)(Figure 4).

### 3.6. eDNA networks reveal previously undescribed relationships

SpIEC-EASI (SParse InversE Covariance estimation for Ecological ASsociation Inference) was used to generate community networks within each marker. We analyzed results tables summarized to family that were filtered for a minimum presence of 15% across the 88 sites, which left 201 18S, 178 16S, and 88 CO1 families. Single marker networks were produced as well as dual marker networks to link 16S with 18S and with CO1, and these were made into interactive figures for users to explore online (e.g. Figure S5, Networks tab). Results produced over 3800 links between families. Two prominent families found to be most integrated in the network were Erythrotrichiaceae (red algae) and Mytilidae (mussels) (Figures S4, Table S12). We queried in the Global Biotic Interactions (GloBI) database, which aggregates published interaction findings, using these two families and 25 others that all had ten or more degrees in the 18S dataset. We asked how many had published biotic interactions, and if so, how many interactions were consistent. Eleven of those families had no reported interactions in GloBI (Table S13). Six of those families had at least one interaction that was detected in our 18S network (Table S12). This result supports the notion that eDNA ecological networks can advance inventories of biotic interactions.

Figure 5 depicts an example of how the interaction networks can be used. The anemone family (Actiniidae) multi-SpIEC-EASI network, determined by 18S and 16S, detected interactions with 13 families. The GloBI query of Actiniidae retrieved 34 interacting families as either sources or targets, but only one family was also found in our results: Symbiodiniaceae. This symbiotic dinoflagellate has been reported elsewhere as part of the anemone holobiome (León-Palmero et al., 2018; Muller et al., 2018). Our networks also revealed a three-way link among Actiniidae, Symbiodiniaceae, and the bacteria Rubritaleaceae. Weber et al. (2017) published the interactions between Rubritaleaceae and Symbiodiniaceae as part of the coral microbiome. These results demonstrate networks depict candidate holobiomes and microbiomes for larger taxa that can further understanding of ecological functions.

**Figure 5.**
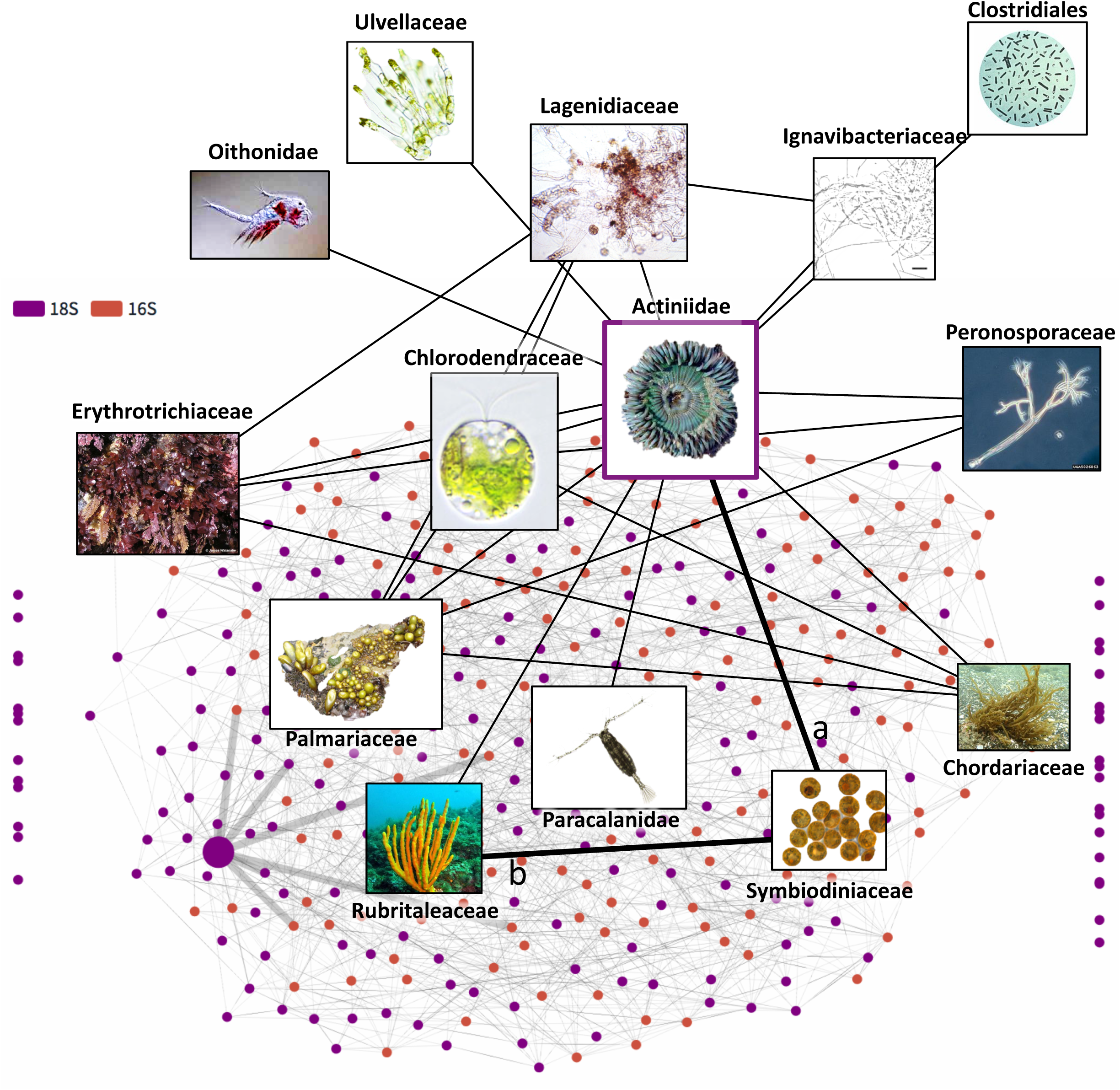
Example network of Actiniidae from the 18S and 16S joint co-occurrence network (background) generated with multiSpiecEasi. a = link previously reported in Muller et al. (2018) and GloBI. b = link previously reported in Weber et al. (2017).

### 3.7. GBIF biodiversity inventories are a different lens of biodiversity

The “Intertidal Biodiversity Survey at Pillar Point” project run by CAS Citizen Science is a focused effort to specifically inventory the low-tide observable biodiversity of the unprotected area tidepools, placing emphasis on tracking certain species such as nudibranchs, and sea stars exposed to sea star wasting disease. Observations in the SMCA or embayment were opportunistic. The survey was sometimes limited when low tide occurred at night. However, the longstanding regularity of citizen science monitoring of Pillar Point (since 2012) has made this beach one of the most iNaturalist observation-rich areas in California. The Survey had 15,539 observations of 596 species on its iNaturalist page when GBIF data were obtained (as of July 5, 2018). Of these observations 12,044 observations were of 457 species and were research grade. We used these data in comparisons with eDNA to understand the differences, parallels, and complementary of the two methods for inventorying and characterizing biodiversity.

The GBIF observations from the sum of the three polygon downloads were largely iNaturalist in origin, comprising 200 of the 226 ‘protected’ (SMCA) polygon records, 1,697 of the 1,799 ‘embayment’ polygon records, and 11,799 of the 11,900 ‘unprotected’ polygon records (Figure 1; Table S2; 13925 records total). Within the GBIF records, we found 689 species were inventoried with iNaturalist, and 140 species were inventoried by other sources. iNaturalist taxa were mostly members of the kingdom Animalia (78%), but also included Chromista (22%), Fungi (1%), and Plantae (17%). The taxonomic lineages identified in GBIF records totaled 598 Animalia, 23 Chromista, 7 Fungi, 147 Plantae, and a single Protozoan, identified as Mycosphaerellaceae (Choanozoa), that was contributed as a natural history museum collection (Tables S2; Figure 1F). Overall, these results highlight the strength of GBIF data to track the animal kingdom (<u>GBIF Sources</u>).

We compared fish in the eDNA 12S and GBIF results because 12S targeted vertebrates. The 12S sequencing results produced a small list, but showed complementarity with GBIF. eDNA 12S results contained 43 taxa, of which 33 were Teleosti and the remainder were other vertebrates (leopard shark, sea otter, sea lion). Of the fish, 23 were resolved to species, with the remainder resolved to genus or family (Table S7). Silversides (Atherinopsidae) were identified in eDNA but sequences were matched to a genus not known in California, Odontesthes. We checked the 12S reference DNA database, and noticed it only contained four Atherinopsidae genera, and the California grunion genus Leuresthes was missing. As of the date of submission of this manuscript, a 12S sequence for this genus has yet to be published in NCBI (April 2019). The grunion genera recorded in GBIF were all missing from the reference eDNA database, suggesting that gaps in sequencing cause erroneous identification, but GBIF information helped elucidate this problem. In a few cases, eDNA could only identify a taxon to genus, such as Xiphister, but of the 71 fish species recorded from Pillar Point in GBIF, there was only a single species in this genus inventoried, the black prickleback Xiphister atropurpureus, providing a candidate source species for the DNA signal. Of the species-level eDNA identifications, twelve species were in GBIF with observations from within the Pillar Point area polygons, and eleven species were in GBIF and observed in California but not observed at Pillar Point (Table S14), suggesting that because intertidal observations are limited to low tides and sight range, GBIF fish observations may represent a more narrow spatiotemporal window than eDNA, and eDNA may fill in gaps of which species visit the reef at periods of inundation.

The most common GBIF observations were families Polyceridae (Mollusca) for Animalia, Laminariaceae (Ochrophyta) for Chromista, Ramalinaceae (Ascomycota) for Fungi, Asteraceae (Thacheophyta) for Plantae. These were not the same as the most frequently observed taxa in eDNA results in those kingdoms: Campanulariidae (Cnidaria) for Animalia, Thraustochytriaceae (Pseudofungi) for Chromista, Glomeraceae (Glomeromycota) for Fungi, and Gigartinaceae (Rhodophyta) for Plantae. These differences in dominant taxa suggest eDNA and GBIF are very different lenses to explore biodiversity. We note that for both datasets, most commonly observed does not equal most common: for photographs, it means more photographs of these taxa have been shared, and for eDNA, it means more samples had DNA from those taxa, and release of DNA into the environment may vary among taxa. The <u>Common Taxa</u> tab shows which taxa are shared between datasets. We were surprised that the Campanulariidae were never identified in GBIF data from the bay, despite their frequency in iNaturalist observations in other parts of California (GBIF.org; not shown). GBIF records did show observations for a taxon in the same order (Leptothecata), but were all for Aglaophenia latirostris. This gap in observations suggests there is capacity to observe broader biodiversity with more overlap with eDNA, but this may require restructuring survey events to generate interest and capacity to notice species.

GBIF and eDNA analyses detected different aspects of biodiversity, with eDNA detecting 82 phyla and GBIF detecting 19 phyla (Figure 1D). The <u>Taxonomy Comparison</u> tab allows users to explore the extent of common taxa at the different phylogenetic levels using Venn diagrams (Figure 1E; Figure S6). With increased taxonomic resolution, the concordance between eDNA and GBIF is diminished (Figure 1D). The family level provided the greatest overlap. Only 48 species-level assignments were common to both eDNA and GBIF. Nonetheless, much could be learned from overlapping taxa (see below).

For example, by comparing frequencies of observation and using the GBIF website (GBIF.org), we were able to develop hypotheses for why the Bay Ghost Shrimp (genus Neotrypaea) was observed more by photographs than with eDNA (Figure 1C). Specifically, we hypothesized that the Bay Ghost Shrimp was more populous or widespread at other times of the year when we did not sample for eDNA (i.e. outside of February and April 2017). GBIF records showed the bulk of observations are in May (n=1701) and June (n=918) while fewer than 25 observations were in each of the other calendar months (Figure S7). This suggests that it may be possible that the early year collections of our eDNA sampling was why we did not observe widespread signals. The shrimp may move to deeper sediments in the winter, or recruit each year. We also observed the inverse pattern for the Star Barnacles (family Chthamalidae; Figure 1C). We detected eDNA signals in all zones, but GBIF data only showed them in the unprotected tidepool zone This may be simply explained by limitations to photograph the adults in the rocky sublittoral zone, or alternatively, that during their larval dispersal phase their planktonic microscopic cyprid larvae are wide spread but are impossible to see in the absence of a plankton tow. Barnacles may also be so common that they are not charismatic to photograph, and this is evidence of human preference. We note that barnacles were not species intentionally surveyed by the volunteers coordinated by CAS.

### 3.8. eDNA fills a gap in GBIF-based biodiversity surveys

GBIF data were too sparse in the SMCA to fairly assess alpha and beta diversity (e.g. Figure 1E). Based on eDNA, we expected that the taxon richness should at least mirror the unprotected polygon. The <u>Area</u> <u>Diversity</u> Table form shows only 103 unique taxa were in the protected SMCA according to GBIF, compared to 520 in the adjacent unprotected tidepools, which is where iNaturalist CAS citizen science activities were concentrated. In contrast, eDNA results limited to the main kingdoms inventoried in GBIF (Animalia, Chromista, and Plantae) had similar taxon richness for adjacent regions: 1210 taxa in the SMCA and 1546 in the unprotected tidepools (see Area Diversity tab).

We then ask if eDNA may help link compositionally similar sites, and chose to examine sites that clustered within the SMCA samples in eDNA beta diversity ordination plots (Figure 2; Figure S8). The site that consistently clustered tightly and with SMCA samples regardless of marker was PP184-B1 (syn K0184-LB-S1; https://data.ucedna.com/samples/792). This single site had ∼100 animal taxa, including cabezon, cormorant, mussels, and ostracods. While it was beyond our capacity to follow replicate GBIF work in the SMCA site and test if these sites are similar using other biodiversity metrics, we propose multi-locus compositional similarity as an eDNA approach to find accessible ‘surrogate sites’ for difficult to access areas.

### 3.9. GBIF taxa and eDNA have numerous singletons

Diminishing overlap with higher taxonomic resolution may be partially explained by misidentifications or rare taxa. Misidentifications can be due to misdiagnosed traits (morphological or DNA sequence). For both eDNA and GBIF, we plotted site frequencies for taxa found, and found datasets were similar across the classification levels for Animalia, the only kingdom with sufficient GBIF data to be fairly comparable (Figure 2). In the eDNA data, 322/742 of species-level taxa were found in only one of the 88 sites (43% singleton species). For GBIF data, 148/566 of species observations were single occurrences (26% singleton species). Although eDNA has significantly more singletons (z-test, z-score −3.6195; p = 0.00015), singletons occur in over a quarter of species-level inventories in the GBIF dataset. This is a substantial proportion that may indicate the same resolution limitations as eDNA. For example, reference databases may be incomplete for both eDNA and GBIF datasets. We also found the large proportion of singletons in the eDNA results were only apparent for Animalia, Chromista, and Plantae, suggesting that in other kingdoms, the eDNA classification bias is not as large. The online <u>Taxonom</u>ic <u>Frequency</u> tab gives users the capacity to study which taxa are rare or common in results.

### 3.10. eDNA detects lower trophic levels than photographic observations

We used the GloBI database to explore biotic of interactions of families detected by GBIF and eDNA observations (Table S13). We compared enrichment in a non-directional GloBI category, ‘interacts_with’, which served as a control, to a directional category, ‘eats’, which served as the test variable. Chi-Square tests (Table 2) showed the eDNA results contained significantly more taxa that are eaten (X-squared = 139.72, df = 1, p-value < 2.2e-16) suggesting that eDNA detects lower trophic levels. No differences were found between eDNA and GBIF results for the “interacts_with” term (p-value >0.8).

**Table 2.**
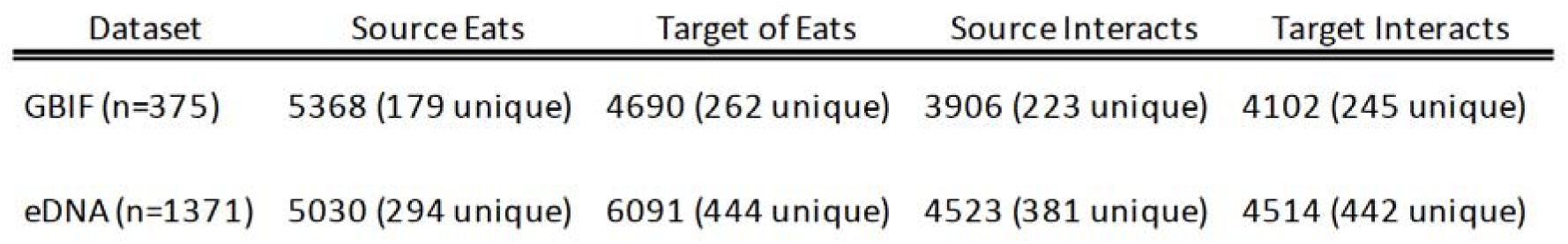
Number of observations of taxa within GBIF and eDNA datasets included in the Global Biotic Interactions database, that were used in Chi-Square tests (see main text).

| Dataset | Source Eats | Target of Eats | Source Interacts | Target Interacts |
| --- | --- | --- | --- | --- |
| GBIF (n=375) | 5368 (179 unique) | 4690 (262 unique) | 3906 (223 unique) | 4102 (245 unique) |
| eDNA (n=1371) | 5030 (294 unique) | 6091 (444 unique) | 4523 (381 unique) | 4514 (442 unique) |

## Discussion

We demonstrate that environmental DNA metabarcoding, an emergent citizen and community science tool, adds rich data on ecology that can also be used in biodiversity assessment. However, eDNA analysis is not likely to infer the same communities as human observation or biological collection surveys. eDNA studies such as ours are likely to capture lower trophic levels than photographic observations (Section 3.10), and depending on the local environment, may capture different spatiotemporal scales (Section 3.1). eDNA may reveal genetic diversity that is as yet unclassifiable and underexplored (Tables S4-S7), and may reveal many novel potential biotic interactions.

Environmental DNA multilocus metabarcoding can also potentially be used to rapidly characterize the spatiotemporal turnover of a system. However, as a recently developed tool, there are important considerations about sensitivity. Relying on eDNA-based estimates of local beta diversity to identify candidate priority conservation areas (da Silva et al., 2018), for instance, is subject to spurious high scores (Section 3.3). Biotic activities, such as growth, spawning, dispersal, burrowing, or recruitment, may lead to misestimation of the spatial ranges of taxa (Section 3.7). Information sourced from GBIF, scientific literature, or taxonomists, along with sufficient DNA barcoding or genome sequencing, is needed to track misidentification in eDNA datasets (Section 3.4).

The metabarcoding approach, which amplifies and sequences barcode loci from a mixed sample (Taberlet et al., 2012; 2018), seeks to match DNA reads to reference sequences from voucher specimens to receive a taxonomic assignment (see Cristescu, 2014 for discussion). DNA barcode reference databases still have large gaps across phylogenies (see Curd et al., 2019) that limit discovery of their true taxonomic membership. Moreover, many barcodes lack diagnostic sequence variants for lower taxonomic assignment (Wolfe et al., 1987). These insufficiencies remain despite decades of generating DNA barcodes (beginning in 1982; Nanney, 1982; CBOL et al., 2009), and the formation of consortia designating specific barcodes (Hebert et al., 2003; Hebert and Gregory, 2005; Yao et al., 2010; Schoch et al., 2012) that have given rise to millions of publicly available sequences (∼6.7 millions sequences in the Barcode of Life Database (BOLD) as of January 11, 2019; boldsystems.org). As natural areas managers rely on species lists and concrete evidence for inventories, we need to close the data gaps while broadening our strategies for effective monitoring from the taxon to the system community where eDNA can be most valuable as tool for ecological research, monitoring and conservation.

Raw observational data from GBIF provide a rich but patchy view of biodiversity in space and time, because observations are not made systematically and they are subject to human bias and the technology they use. We are limited by where people can go and by their observational choices and capacities to take photographs or make collections (Section 3.8). The CAS team is currently developing tools to manage these biases. Haphazard sampling for eDNA can have the same bias, but with little effort, systematic sampling events can be organized and executed by volunteers. Accurate identification is also a potential challenge for iNaturalist as it is for eDNA. For photographic observations, machine learning algorithms assist the community in identifications, but the computer vision model is limited to 10,000 species currently (https://www.inaturalist.org/pages/computer_vision_demo). Community experts affirm and revise these identifications, but over half of the species identified in iNaturalist have fewer than 20 total observations, as a result, there is little comparative data available to corroborate taxonomic assignments. We found elevated single occurrences of taxa in both eDNA and GBIF datasets, which may partially be explained by misidentification (Section 3.9; Figure 6). The addition of more data will be necessary to distinguish rare from misidentified species, as will integration of different kinds of biodiversity inventories in community platforms such as GBIF. Because of the accessibility of various data sources on the GBIF platform, we were able to show 11 fish species found with eDNA are first reports at the beach, as were 76 phyla that had no record in GBIF, which showcases eDNA as a feasible method to both fill inventory gaps and to include the more challenging branches of the tree of life (Ruppert et al., 2019).

**Figure 6.**
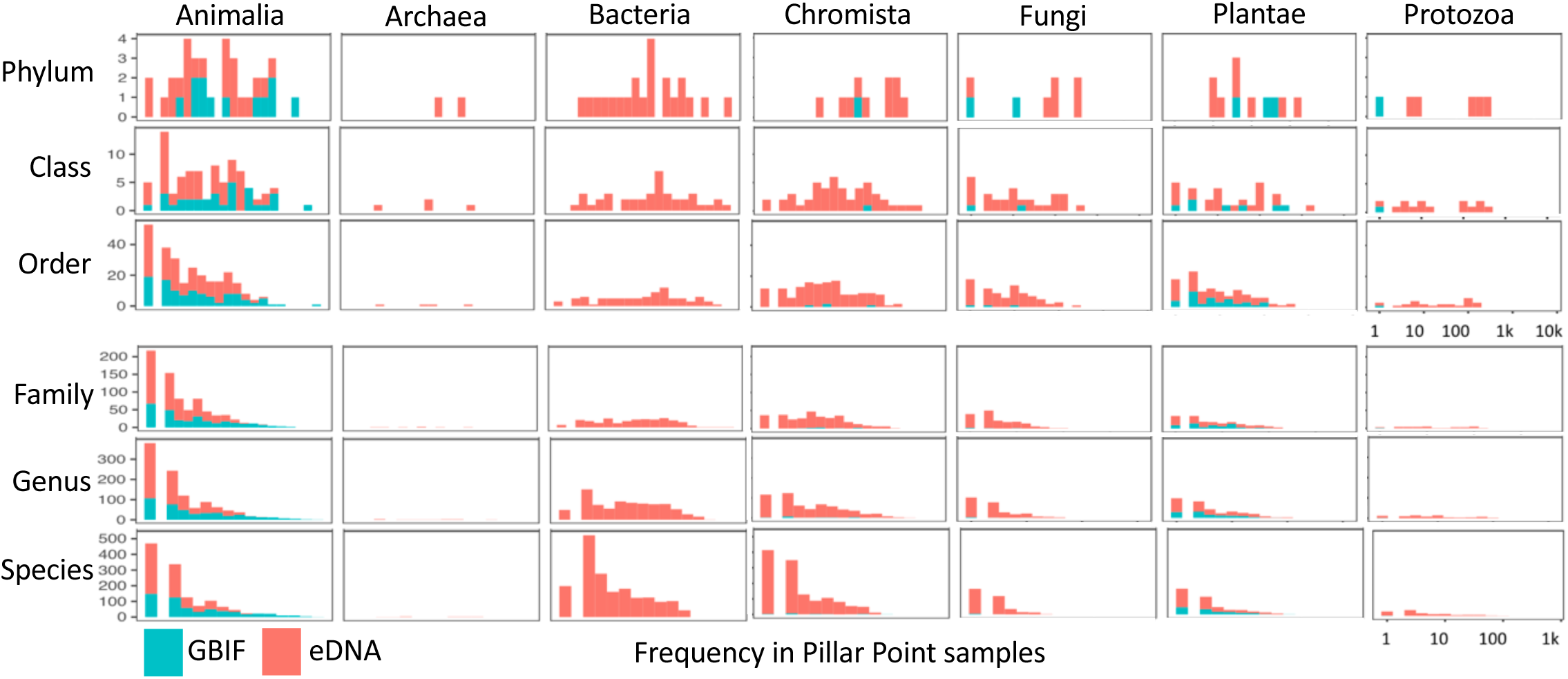
Log_10_ transformed plot of taxon frequency observed in eDNA or GBIF datasets. All taxa were used if they had been classified minimally to the classification level designated (y-axis). The x-axis is the number of sites where a taxon was observed. Most observations in GBIF are Animalia. Single observations are high for both eDNA and GBIF at the levels of families through species, most pronounced in Animalia.

The pace of change in the science of environmental DNA sequencing and analysis is rapid, which makes the value of environmental sample collections all the more important. Curatorial resources such as the Global Genome Biodiversity Network, the NHMLA Diversity Initiative for the Southern California Ocean (DISCO), and the Genomic Observatories Metadatabase (GeOMe), are helping samples and their associated metadata remain accessible for future research. As techniques for sequencing and analyzing eDNA improve, we may soon readily track intraspecific change or genetic diversity that relates to ecological functions. Projects such as for the Earth BioGenome Project (Lewin et al., 2018) may also use environmental samples for species discovery critical to fill in the tree of life and enhance reference databases. Our brief investigation into Ostracoda genetic variants suggests surveying the unprotected tidepools of Pillar Point are the most likely to have uncataloged species (Appendix 2; Table S8). The value of eDNA-based ecological community networks (Section 3.6; Figure S5), such as the candidate holobiomes (Figure 5) they reveal, may also increase as technology and theory improve.

The Pillar Point marsh and inner beach had been graded C-F in the most recent Heal the Bay report card (2016). While eDNA did show high density of harmful pathogens (Figure 4), we also find a complex, connected, and extremely diverse environment in both the marsh (highest alpha diversity; dense with complex life cycle parasites) and inner beach (highest LCBD). Our demonstration of eDNA to catalog the rich and dynamic biodiversity of this beach provides exciting evidence that there are many ‘low hanging fruit’ findings pertinent to both management and to basic research in biodiversity that can be gleaned from a few days of fieldwork and <$4000 USD in DNA library preparation and sequencing costs.

One of the major conservation questions arising from this case study unique about concerned defining and monitoring the unique characteristics of the SMCA. With eDNA, the SMCA was found to harbor high density of unmonitored groups such as urochordates (Ascidiacea; with known invasive species), marine flatworms (Polycladida), and a unique assemblage of microbial eukaryotes. However, a dearth of photographic data contributions meant many priority taxa were not tracked. Although enhancing the observational data for the SMCA may be possible, our results at least provide a framework for monitoring the system for perturbation without relying on human observation. We suggest that by using surrogate sites identified by low eDNA beta diversity (Section 3.8), responses to environmental stress or change can be measured in heavily surveyed sites, and these can be used to predict the processes occurring in the SMCA.

The CALeDNA Program is growing its online inventory of California’s biodiversity and participants in the program are encouraged to pair their eDNA sampling with iNaturalist observations. Future directions will include more case studies that compare both tools with additional biodiversity correlates such as those from remote sensing data. We will also work toward better understanding the significance of spatiotemporal trends. It is imperative that CALeDNA and various public and professional biomonitoring programs work together to cross-inform each other so we can make strategic efforts to promote healthy biodiversity.

## Acknowledgements

Support for this project was provided by the University of California Catalyst Program CA-16-376437. We thank the volunteers who collected CALeDNA samples and iNaturalist observations. We thank M. Delaney of NOAA and thank the California Dept. of Fish and Wildlife for assistance with permits and permissions. We thank M. Lin, and Z. Gold for assistance with analyses. The authors (AW, NDP, RW) graciously acknowledge NHM Marine Biodiversity Center infrastructure support from colleagues, Kathy Omura and Jenessa Wall. This is Contribution Number 5 of the NHM Diversity Initiative of the Southern California Ocean.

## Data Availability

All raw sequence data are available in the NCBI SRA under project PENDING ACCEPTANCE. Software necessary to repeat the analyses are available in DRYAD (https://doi.org/10.5061/dryad.mf0126f), which also contains DNA reference databases for 18S, 12S, and 16S markers. The CO1-BOLD reference database is archived in Zenodo (PENDING). All other data needed to replicate the results are included in supplemental materials.

## Legends for Supplementary Figures, Supplementary Tables

**Figure S1.** Taxon accumulation curves as increasing reads are retained in rarified datasets. Curves were used to choose rarefaction settings.

**Figure S2.** Alpha diversity plots calculated using species richness, Simpson’s index, and the Shannon index (H’) in a. Only Shannon index is shown in b.

**Figure S3.** Relative abundance barplots coloring the top 21 species. Grouping definitions are in Table S1 and Figure S2.

**Figure S4.** Plots to interpret skewness of LCBD scores if holobiomes of DNA swamping taxa are left in or removed from 18S results. The dependent variables are binary for either swamping or significantly different LCBD scores from the rest of their groups. The dependent variables are different representations of the LCBD score range. A. Plot showing the samples with significant LCBD P-values in original results experienced either an increase in LCBD or a decrease in LCBD when holobiome-reduced results were used to recalculate LCBD scores. B. Plot showing the majority of ‘swamped’ samples experienced a reduction in LCBD when holobiome-reduced results were used to recalculate LCBD. C. Plot showing the samples that were originally ‘swamped’ now fall into the distribution range of the bulk of the samples in the holobiome-reduced LCBD results. D. Plot showing the samples that were originally ‘swamped’ were distributed above the LCBD of the bulk of the samples in the original LCBD results. Comparing plots D and C demonstrates reduction of the candidate holobiomes of high abundance taxa helped the samples that experienced swamping look typical.

**Figure S5.** SpiecEasi network analysis results of families detected by eDNA. Top left is a density plot of the number of degrees taxa have in each network type. SpiecEasi networks are shown with numbers corresponding to a family with the key in Table S12. Interactive figures of the networks are available on the Pillar Point website.

**Figure S6.** Degree of overlapping taxa among polygons. Circle sizes are scaled to the number of taxa.

**Figure S7.** Excerpt from the GBIF website of observation frequency over months of the year.

**Figure S8.** Ordination plots as in Figure 2 but with sample names as labels. PP184-B1 is the unprotected tidepool sample that is closest to the SMCA samples in both 16S and 18S datasets.

**Table S1.** eDNA sample metadata

**Table S2.** GBIF data for the three polygons

**Table S3.** Comparison of CO1 settings

**Table S4.** eDNA decontaminated results for 16S marker

**Table S5.** eDNA decontaminated results for the 18S marker

**Table S6.** Raw Anacapa output from MiSeq runs and eDNA decontaminated results for the CO1 marker

**Table S7.** eDNA decontaminated results for the 12S marker

**Table S8.** Genetic diversity analysis of eDNA results for Ostracods

**Table S9.** ANOSIM results from Raup and Jaccard beta diversity analyses

**Table S10.** Local Contribution to Beta Diversity scores

**Table S11.** Density enrichment test results and statistics

**Table S12.** Family co-occurrence networks

**Table S13.** Global Biotic Interactions data and results

## References

Allen, W. (2003). Plant blindness. BioScience 53, 926–926.

Amaral-Zettler, L.A., McCliment, E.A., Ducklow, H.W., and Huse, S.M. (2009). A method for studying protistan diversity using massively parallel sequencing of v9 hypervariable regions of small-subunit ribosomal RNA Genes. PLOS ONE doi:10.1371/journal.pone.0006372.

Auld, S.K., and Tinsley, M.C. (2015). The evolutionary ecology of complex lifecycle parasites: linking phenomena with mechanisms. Heredity 114, 125–132.

Benjamini, Y., and Hochberg, Y. (1995). Controlling the false discovery rate: a practical and powerful approach to multiple testing. Journal of the Royal Statistical Society Series B 57, 289–300.

Bird, T.J., Bates, A.E., Lefcheck, J.S., Hill, N.A., Thomson, R.J., Edgar, G.J., Stuart-Smith, R.D., Wotherspoon, S., Krkosek, M., Stuart-Smith, J.F., et al. (2014). Statistical solutions for error and bias in global citizen science datasets. Biological Conservation 173, 144–154.

Bradley, I.M., Pinto, A.J., and Guest, J.S. (2016). Design and Evaluation of Illumina MiSeq-Compatible, 18S rRNA Gene-Specific primers for improved characterization of mixed phototrophic communities. Applied Environmental Microbiology 82, 5878–5891.

Callahan, B.J., McMurdie, P.J., Rosen, M.J., Han, A.W., Johnson, A.J.A., and Holmes, S.P. (2016). DADA2: High-resolution sample inference from Illumina amplicon data. Nature Methods 13, 581–583.

Caporaso, J.G., Lauber, C.L., Walters, W.A., Berg-Lyons, D., Huntley, J., Fierer, N., Owens, S.M., Betley, J., Fraser, L., Bauer, M., et al. (2012). Ultra-high-throughput microbial community analysis on the Illumina HiSeq and MiSeq platforms. The ISME Journal 6, 1621–1624.

CBOL Plant Working Group, Hollingsworth, P.M., Forrest, L.L. et al. (2009). A DNA barcode for land plants. Proceedings of the National Academy of Sciences 106,12794–12797. doi:10.1073/pnas.0905845106.

Chase, J.M., Kraft, N.J.B., Smith, K.G., Vellend, M., and Inouye, B.D. (2011). Using null models to disentangle variation in community dissimilarity from variation in α-diversity. Ecosphere. doi:10.1890/ES10-00117.1.

Cristescu, M.E. (2014). From barcoding single individuals to metabarcoding biological communities: towards an integrative approach to the study of global biodiversity. Trends in Ecology and Evolution 29, 566–571.

Curd, E.E., Gold, Z., Kandlikar, G., Gomer, J., Ogden, M., O’Connell, T., Pipes, L., Schweizer, T., Rabichow, L., Lin, M., et al. (2019). Anacapa Toolkit: an environmental DNA toolkit for processing multilocus metabarcode datasets. Methods in Ecology and Evolution, accepted and in press. BioRxiv doi:10.1101/488627.

Csardi, G., Nepusz, T. (2006). The Igraph software package for complex network research. InterJournal Complex Systems, 1695.

da Silva, P.G., Hernández, M.I.M., and Heino, J. (2018). Disentangling the correlates of species and site contributions to beta diversity in dung beetle assemblages. Diversity and Distributions 24, 1674–1686.

Davis, N.M., Proctor, D.M., Holmes, S.P., Relman, D.A., and Callahan, B.J. (2018). Simple statistical identification and removal of contaminant sequences in marker-gene and metagenomics data. Microbiome 6, 226.

Diaz, S., Settele, J., Brondizio, E. (2019). Summary for policymakers of the global assessment report on biodiversity and ecosystem services of the Intergovernmental Science-Policy Platform on Biodiversity and Ecosystem Services. Germany: Secretariat of the Intergovernmental SciencePolicy Platform on Biodiversity and Ecosystem Services. Available from https://www.ipbes.net/sites/default/files/downloads/spm_unedited_advance_for_posting_htn.pdf

Dray, S., Blanchet, G., Borcard, D. et al. (2016). adespatial: Multivariate multiscale spatial analysis. Retrieved from https://cran.r-project.org/web/packages/adespatial/index.html.

Dunn, O.J. (1964). Multiple comparisons using rank sums. Technometrics 6, 241–252.

Folmer, O., Black, M., Hoeh, W., Lutz, R., and Vrijenhoek, R. (1994). DNA primers for amplification of mitochondrial cytochrome c oxidase subunit I from diverse metazoan invertebrates. Mol Mar Biol Biotechnol 3, 294–9.

Fonseca, V.G., Carvalho, G.R., Sung, W., Johnson, H.F., Power, D.M., Neill, S.P., Packer, M., Blaxter, M.L., Lambshead, P.J.D., Thomas, W.K., et al. (2010). Second-generation environmental sequencing unmasks marine metazoan biodiversity. Nature Communications. doi:10.1038/ncomms1095.

Gao, X., Lin, H., Revanna, K., and Dong, Q. (2017). A Bayesian taxonomic classification method for 16S rRNA gene sequences with improved species-level accuracy. BMC Bioinformatics. doi:10.1186/s12859-017-1670-4.

Groom, Q., Weatherdon, L., and Geijzendorffer, I.R. (2017). Is citizen science an open science in the case of biodiversity observations? Journal of Applied Ecology 54, 612–617.

Guerrero, R., Margulis, L., and Berlanga, M. (2013). Symbiogenesis: the holobiont as a unit of evolution. International Microbiology 133–143.

Hebert Paul D. N., Cywinska Alina, Ball Shelley L., and deWaard Jeremy R. (2003). Biological identifications through DNA barcodes. Proceedings of the Royal Society of London. Series B: Biological Sciences 270, 313–321.

Hebert, P.D.N., and Gregory, T.R. (2005). The promise of DNA barcoding for taxonomy. Systematic Biology 54, 852–859.

Hecker, S., Bonney, R., Haklay, M., Hölker, F., Hofer, H., Goebel, C., Gold, M., Makuch, Z., Ponti, M., Richter, A., et al. (2018). Innovation in Citizen Science – Perspectives on Science-Policy Advances. Citizen Science: Theory and Practice 3, 4.

Hechinger, R.F., and Lafferty, K.D. (2005). Host diversity begets parasite diversity: bird final hosts and trematodes in snail intermediate hosts. Proceedings of the Royal Society B: Biological Sciences 272, 1059–1066.

Herlemann, D.P.R., Lundin, D., Labrenz, M., Jürgens, K., Zheng, Z., Aspeborg, H., and Andersson, A.F. (2013). Metagenomic de novo assembly of an aquatic representative of the Verrucomicrobial class Spartobacteria. MBio 4, e00569–12.

Hollander, M., Wolfe, D.A., and Chicken, E. (2013). Nonparametric Statistical Methods USA: John Wiley and Sons, Inc.

Hupało, K., Mamos, T., Wrzesińska, W., and Grabowski, M. (2018). First endemic freshwater Gammarus from Crete and its evolutionary history—an integrative taxonomy approach. PeerJ 6.

Kandlikar, G.S., Gold, Z.J., Cowen, M.C., Meyer, R.S., Freise, A.C., Kraft, N.J.B., Moberg-Parker, J., Sprague, J., Kushner, D.J., and Curd, E.E. (2018). ranacapa: An R package and Shiny web app to explore environmental DNA data with exploratory statistics and interactive visualizations. F1000Research. doi:10.12688/f1000research.16680.1.

Keeley, J. E., and P. H. Zedler. (1998). Characterization and global distribution of vernal pools. Pages 1–14 in C. W. Witham, E. T. Bauder, D. Belk, W. R. Ferren Jr., and R. Ornduff. editors. Ecology, conservation and management of vernal pool ecosystems–Proceedings from a 1996 conference. California Native Plant Society, Sacramento.

Kurtz, Z.D., Müller, C.L., Miraldi, E.R., Littman, D.R., Blaser, M.J., and Bonneau, R.A. (2015). Sparse and compositionally robust inference of microbial ecological networks. PLOS Computational Biology. doi:10.1371/journal.pcbi.1004226.

Langmead, B., and Salzberg, S.L. (2012). Fast gapped-read alignment with Bowtie 2. Nature Methods 9, 357–359.

Legendre, P., and Cáceres, M.D. (2013). Beta diversity as the variance of community data: dissimilarity coefficients and partitioning. Ecology Letters 16, 951–963.

León-Palmero, E., Joglar, V., Álvarez, P.A., Martín-Platero, A., Llamas, I., and Reche, I. (2018). Diversity and antimicrobial potential in sea anemone and holothurian microbiomes. PLOS. doi:10.1371/journal.pone.0196178.

Leray, M., Yang, J.Y., Meyer, C.P., Mills, S.C., Agudelo, N., Ranwez, V., Boehm, J.T., and Machida, R.J. (2013). A new versatile primer set targeting a short fragment of the mitochondrial COI region for metabarcoding metazoan diversity: application for characterizing coral reef fish gut contents. Frontiers in Zoology. doi:10.1186/1742-9994-10-34.

Lewin, H.A., Robinson, G.E., Kress, W.J., Baker, W.J., Coddington, J., Crandall, K.A., Durbin, R., Edwards, S.V., Forest, F., Gilbert, M.T.P., et al. (2018). Earth BioGenome Project: Sequencing life for the future of life. PNAS 115, 4325–4333.

Martin, M. (2011). Cutadapt removes adapter sequences from high-throughput sequencing reads. EMBnet.Journal 17, 10–12.

McMurdie, P.J., and Holmes, S. (2013). phyloseq: An R package for reproducible interactive analysis and graphics of microbiome census data. PLOS ONE. doi:10.1371/journal.pone.0061217.

Meyer, R.S., Curd, E.E., Schweizer, T., Gold, Z., Ruiz Ramos, D., Shirazi, S., Kandlikar, G., Kwan, W.-Y., Lin, M., Friese, A., et al. (2019). The California environmental DNA “CALeDNA” program. BioRxiv. doi:10.1101/503383.

Meyer, R., and Drill, S. (2019). Community and citizen science at the UC Division of Agriculture and Natural Resources. Final Report, Center for Community and Citizen Science. https://education.ucdavis.edu/sites/main/files/citizen_and_community_science_at_anr_final_report.pdf

Miya, M., Sato, Y., Fukunaga, T., Sado, T., Poulsen, J.Y., Sato, K., Minamoto, T., Yamamoto, S., Yamanaka, H., Araki, H., et al. (2015). MiFish, a set of universal PCR primers for metabarcoding environmental DNA from fishes: detection of more than 230 subtropical marine species. Royal Society Open Science. doi:10.1098/rsos.150088.

Muller, E.M., Fine, M., and Ritchie, K.B. (2016). The stable microbiome of inter and sub-tidal anemone species under increasing pCO_2_. Scientific Reports 6, 37387. doi:10.1038/srep37387.

Nanney, D.L. (1982). Genes and phenes in Tetrahymena. BioScience 32, 783–788.

National Research Council. (2001). Grand challenges in environmental sciences. National Research Council. doi: 10.17226/9975

O’Donnell, J.L., Kelly, R.P., Shelton, A.O., Samhouri, J.F., Lowell, N.C., and Williams, G.D. (2017). Spatial distribution of environmental DNA in a nearshore marine habitat. PeerJ. doi:10.7717/peerj.3044.

Oksanen, J., Blanchet, F., Kindt, R. (2015). Community ecology package ‘vegan’. Retrieved from https://github.com/vegandevs/vegan.

Patterson, D., Mozzherin, D., Shorthouse, D., and Thessen, A. (2016). Challenges with using names to link digital biodiversity information. Biodiversity Data Journal 4. doi:10.3897/BDJ.4.e8080.

Peters, M.K., Hemp, A., Appelhans, T., Becker, J.N., Behler, C., Classen, A., et al. (2019). Climate–land-use interactions shape tropical mountain biodiversity and ecosystem functions. Nature 568, 88–92.

Poelen, J.H., Simons, J.D., and Mungall, C.J. (2014). Global biotic interactions: An open infrastructure to share and analyze species-interaction datasets. Ecological Informatics 24, 148–159.

Raup, D.M., and Crick, R.E. (1979). Measurement of faunal similarity in paleontology. Journal of Paleontology 53, 1213–1227.

Robertson, T., Döring, M., Guralnick, R., Bloom, D., Wieczorek, J., Braak, K., Otegui, J., Russell, L., and Desmet, P. (2014). The GBIF integrated publishing toolkit: facilitating the efficient publishing of biodiversity data on the internet. PLOS ONE. doi:10.1371/journal.pone.0102623.

Ruggiero, M.A., Gordon, D.P., Orrell, T.M., Bailly, N., Bourgoin, T., Brusca, R.C., Cavalier-Smith, T., Guiry, M.D., and Kirk, P.M. (2015). A higher level classification of all living organisms. PLOS ONE 10. doi:10.1371/journal.pone.0119248.

Ruppert, K.M., Kline, R.J., and Rahman, M.S. (2019). Past, present, and future perspectives of environmental DNA (eDNA) metabarcoding: A systematic review in methods, monitoring, and applications of global eDNA. Global Ecology and Conservation 17, e00547.

Schoch, C.L., Seifert, K.A., Huhndorf, S., Robert, V., Spouge, J.L., Levesque, C.A., Chen, W., and Consortium, F.B. (2012). Nuclear ribosomal internal transcribed spacer (ITS) region as a universal DNA barcode marker for Fungi. PNAS 109, 6241–6246.

Taberlet, P., Bonin, A., Zinger, L., and Coissac, E. (2018). Environmental DNA: For Biodiversity Research and Monitoring. United Kingdom: Oxford University Press.

Taberlet P, Coissac E, Pompanon F et al. 2012. Towards next-generation biodiversity assessment using DNA metabarcoding. Molecular Ecology, 21:2045–2050. doi:10.1111/j.1365-294X.2012.05470.x.

Tipton, L., Müller, C.L., Kurtz, Z.D., Huang, L., Kleerup, E., Morris, A., Bonneau, R., and Ghedin, E. (2018). Fungi stabilize connectivity in the lung and skin microbial ecosystems. Microbiome. doi:10.1186/s40168-017-0393-0.

Weber, L., DeForce, E., and Apprill, A. (2017). Optimization of DNA extraction for advancing coral microbiota investigations. Microbiome. doi:10.1186/s40168-017-0229-y.

Whittaker, R.H. (1972). Evolution and measurement of species diversity. Taxon 21, 213–251.

Wickham, H. (2009) ggplot2: elegant graphics for data analysis. USA: Springer.

Wolfe, K.H., Li, W.H., and Sharp, P.M. (1987). Rates of nucleotide substitution vary greatly among plant mitochondrial, chloroplast, and nuclear DNAs. Proceedings of the National Academy of Sciences 84, 9054–9058.

Yao, H., Song, J., Liu, C., Luo, K., Han, J., Li, Y., Pang, X., Xu, H., Zhu, Y., Xiao, P., et al. (2010). Use of ITS2 region as the universal DNA barcode for plants and animals. PLOS ONE. doi:10.1371/journal.pone.0013102.

Zinger, L., Taberlet, P., Schimann, H., Bonin, A., Boyer, F., Barba, M.D., Gaucher, P., Gielly, L., Giguet-Covex, C., Iribar, A., et al. (2019a). Body size determines soil community assembly in a tropical forest. Molecular Ecology 28, 528–543.

Zinger, L., Bonin, A., Alsos, I.G., Bálint, M., Bik, H., Boyer, F., Chariton, A.A., Creer, S., Coissac, E., Deagle, B.E., et al. (2019b). DNA metabarcoding—Need for robust experimental designs to draw sound ecological conclusions. Molecular Ecology 28, 1857–1862.

