## Supplementary material for "Beach environmental DNA fills gaps in photographic biomonitoring to track spatiotemporal community turnover across 82 phyla": Table 1

| Website Tab | Function |
| --- | --- |
| **Intro** | Introduces the CALeDNA and GBIF data, and shows the location of the data points on a map of Pillar Point |
| **Occurrence Comparison** | Shows the distribution of total presence counts of a taxon in eDNA results or in GBIF observations. |
| **GBIF Sources** | Depicts data distribution from iNaturalist vs other contributions |
| **GBIF Taxonomy** | Shows the GBIF taxa, whether they are in the NCBI database, and whether they have an eDNA result that matches |
| **Common Taxa** | Lists taxa at each classification level that overlap, showing datapoints on the Pillar Point map |
| **Area Diversity** | Presents filterable comparisons of unique and common taxa found among the three polygons for eDNA and GBIF dataset |
| **Taxonomy Comparison** | Displays Venn diagrams of filterable unique and common taxa between eDNA and GBIF results |
| **Detection Frequency** | Presents a filterable sorted list of frequency of occurrences for all taxa in a classification level |
| **Networks** | Displays interactive co-occurrence networks at the family level |
| **Biotic Interactions** | Presents species frequently observed at Pillar Point with iNaturalist, the known biotic interactions and whether interacting taxa were in GBIF or eDNA datasets |
