## Supplementary figures and images for "Beach environmental DNA fills gaps in photographic biomonitoring to track spatiotemporal community turnover across 82 phyla"

### Figure S1

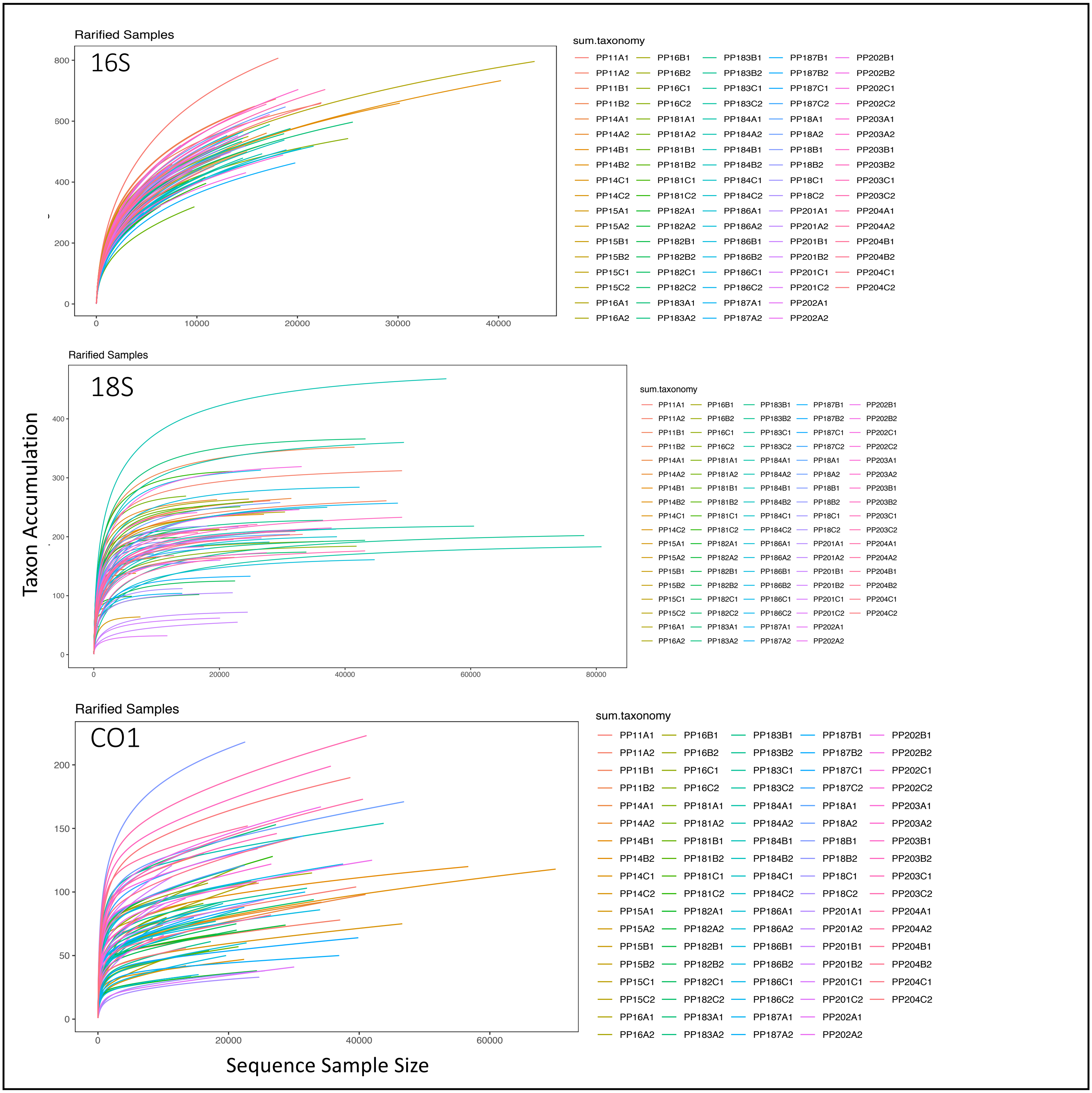

### Figure S2

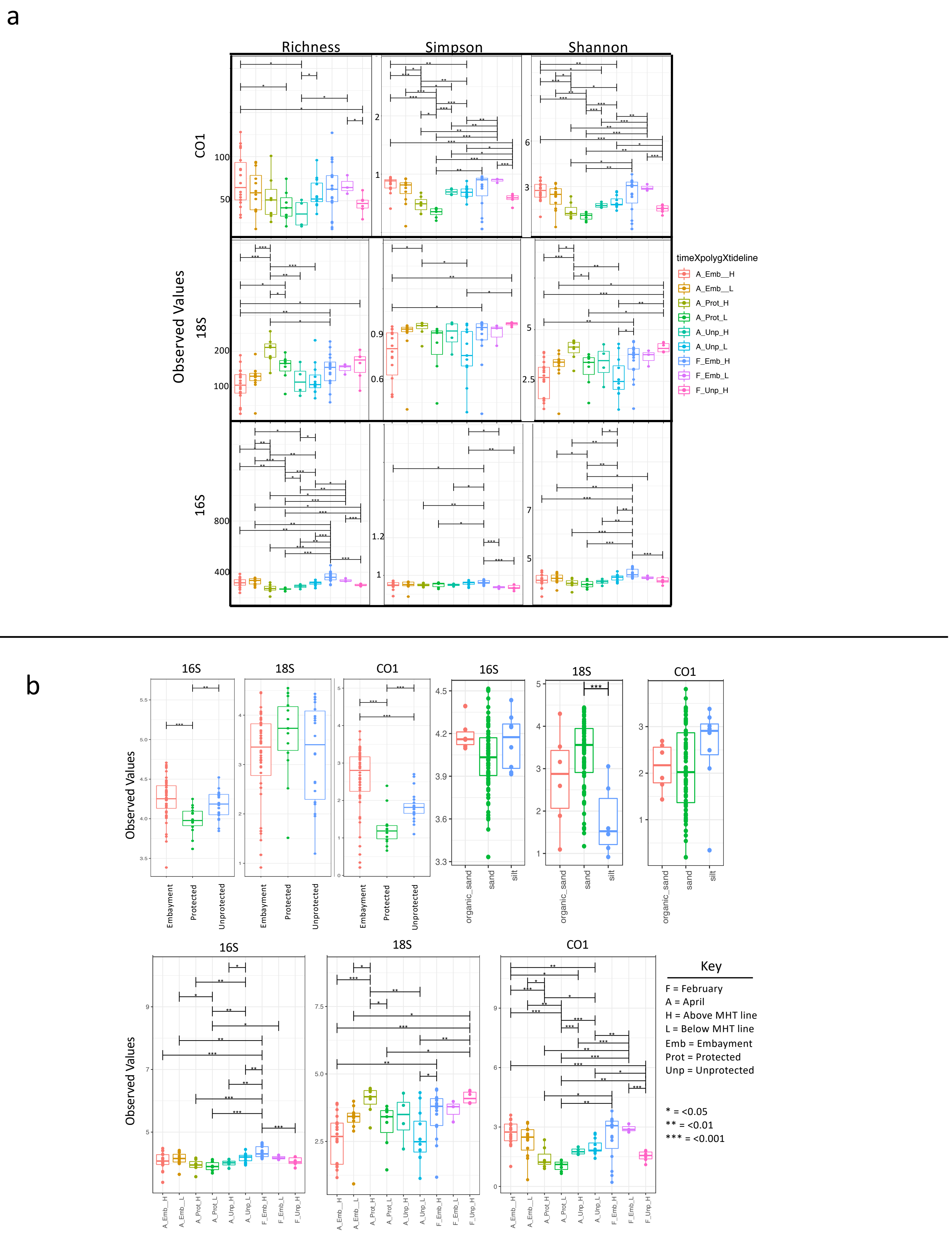

### Figure S3

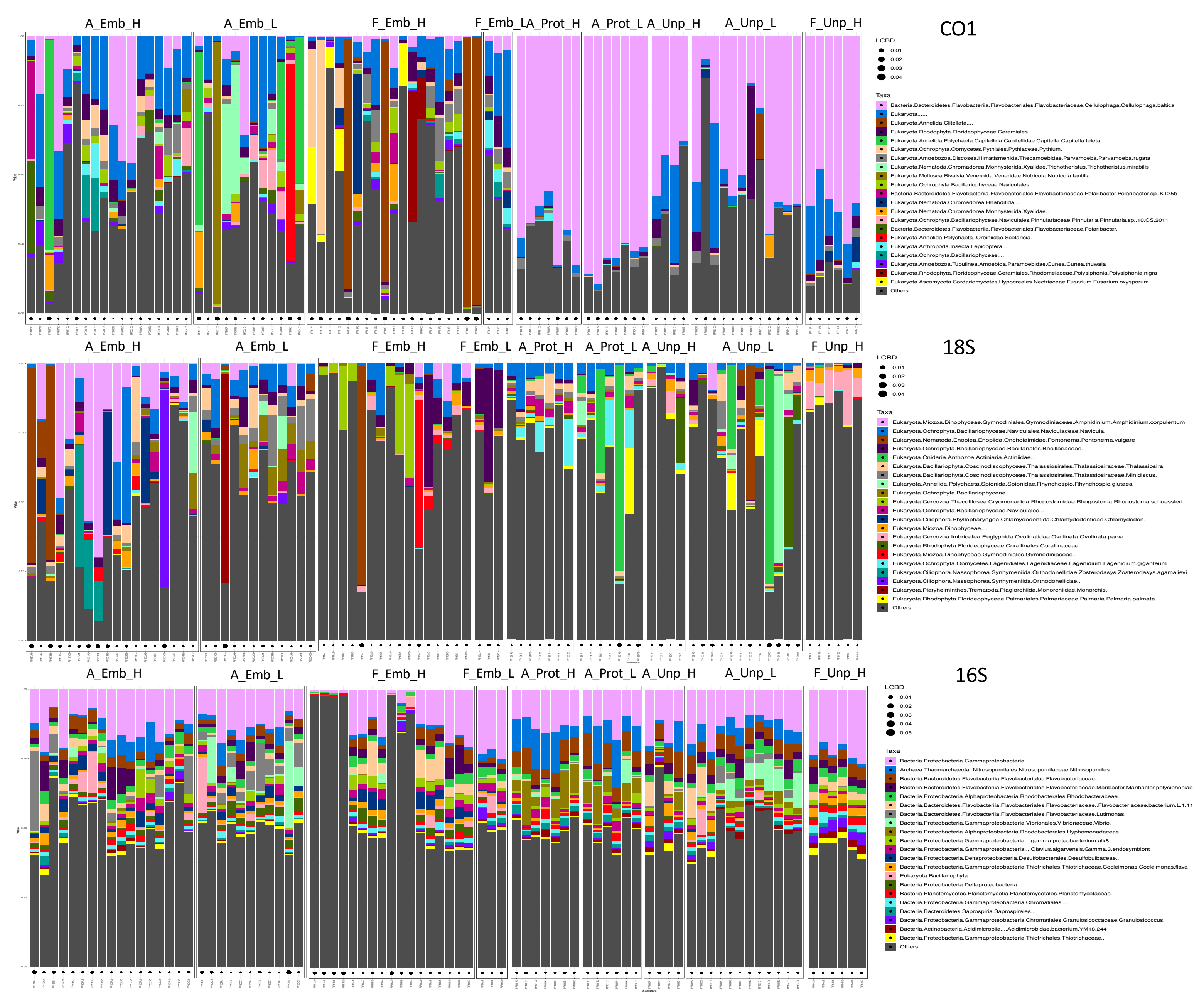

### Figure S4

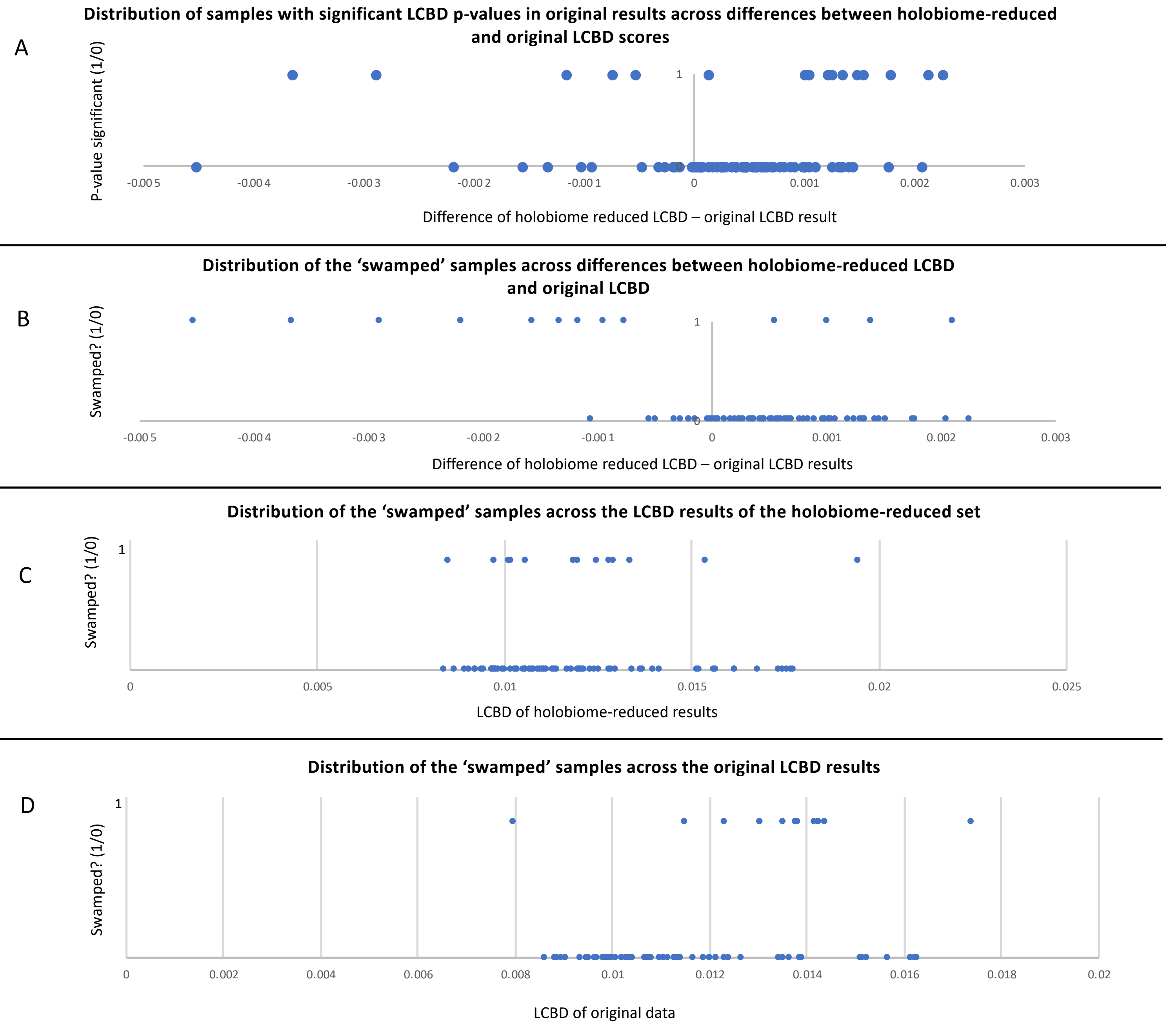

### Figure S5

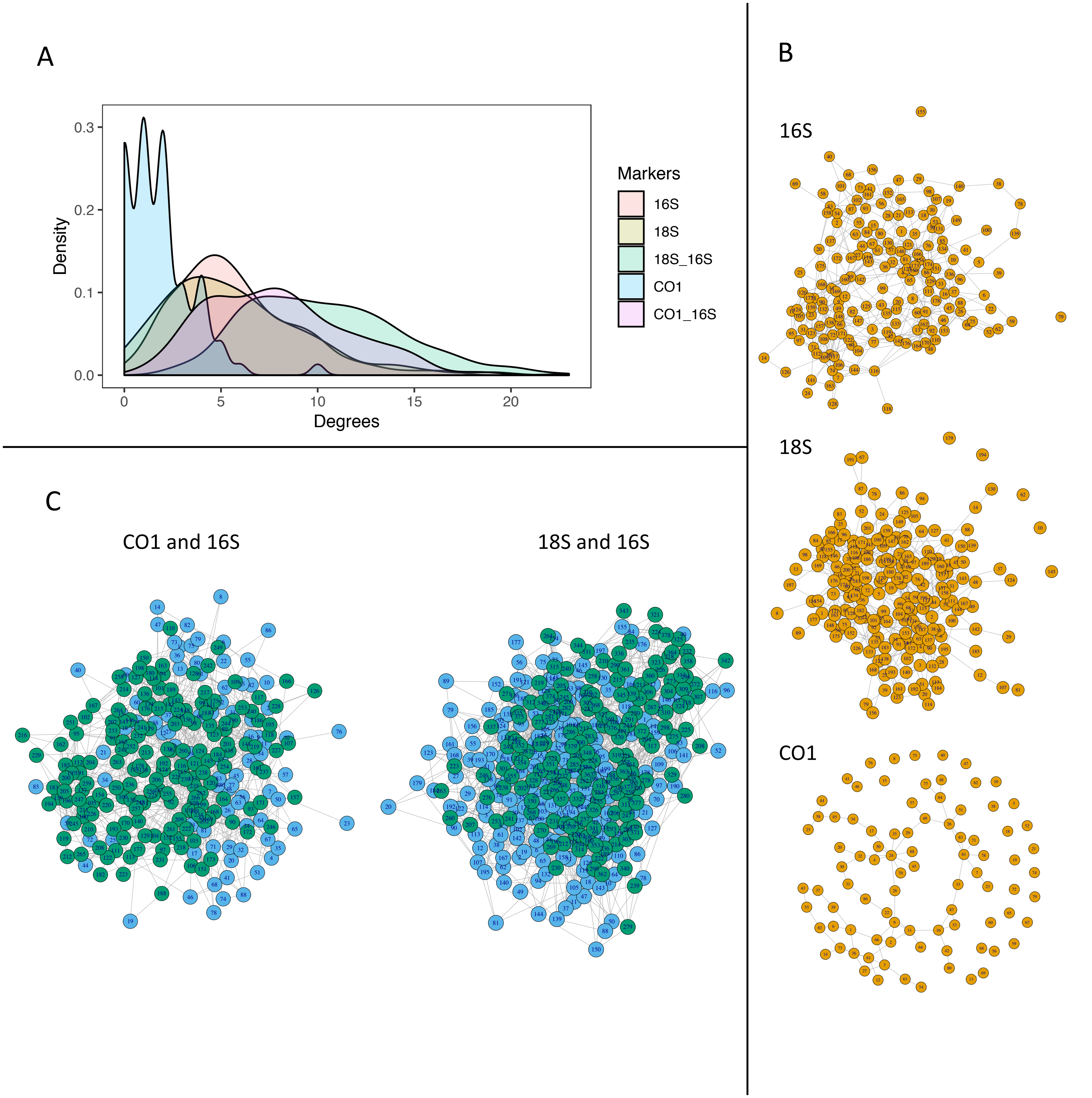

### Figure S6

| Rank | GBIF/eDNA/overlap | GBIF/eDNA/overlap |
|---|---|---|
| Phylum | 19/76/19 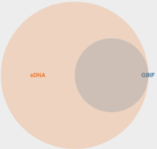  | 12/23/12 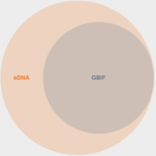  |
| Class | 43/213/40 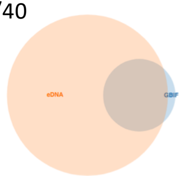  | 32/55/29 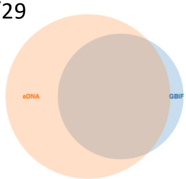  |
| Order | 156/591/98 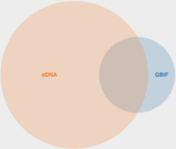  | 109/163/61 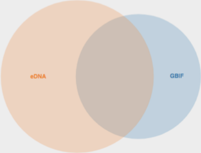  |
| Family | 357/1370/115   | 282/477/76 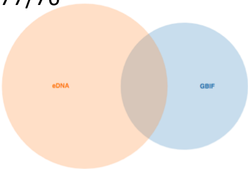  |
| Genus | 563/2656/79 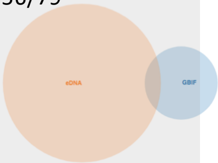  | 424/667/45 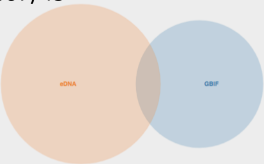  |
| Species | 726/4602/35 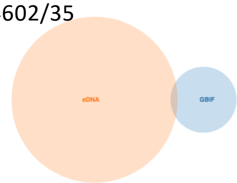  | 566/742/22 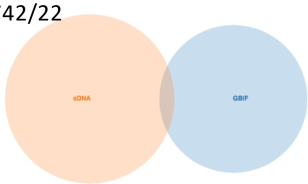  |

### Figure S8

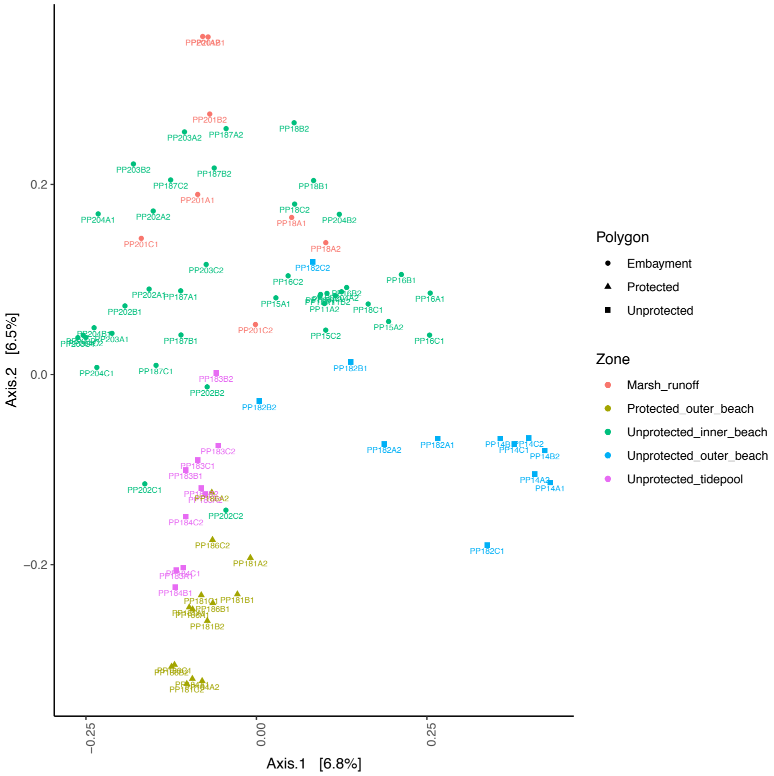

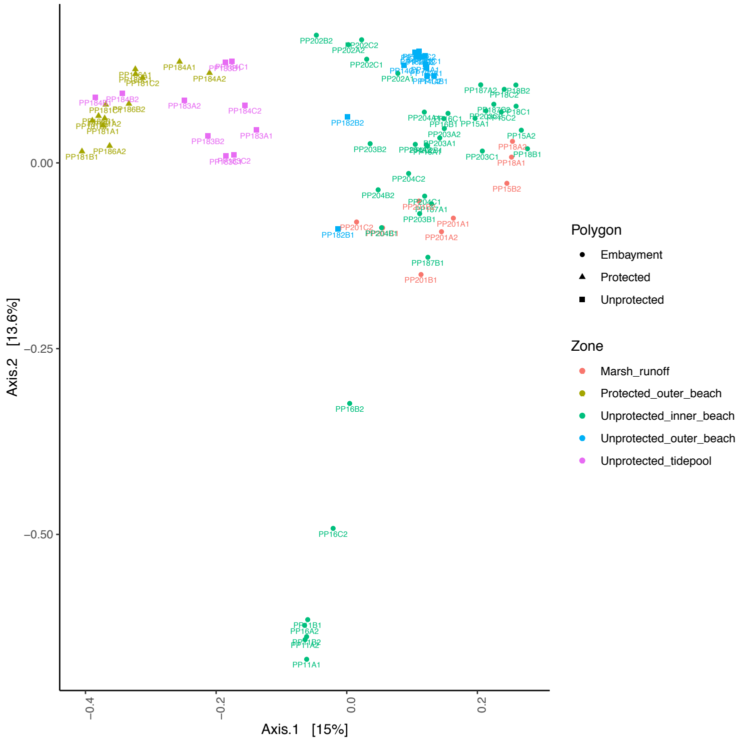
