## Supplementary material for "Beach environmental DNA fills gaps in photographic biomonitoring to track spatiotemporal community turnover across 82 phyla": Figure S7

SPECIES | ACCEPTED

### *Neotrypaea californiensis* (Dana, 1854)

Published in: Dana, J.D. (1854). Catalogue and descriptions of Crustacea collected in California by Dr. John L. Le Conte. *Proceedings of the Academy of Natural Sciences of Philadelphia*. 7: 175-177.
source: Catalogue of Life

In: GBIF Backbone Taxonomy

**Bay ghost shrimp** In English **Basionym:** *Callianassa californiensis* Dana, 1854



#### OCCURRENCES PER MONTH

| Month | Count |
|---|---|
| January | 9 |
| February | 8 |
| March | 13 |
| April | 14 |
| May | 1,701 |
| June | 918 |
| July | 25 |
| August | 22 |
| September | 9 |
| October | 2 |
| November | 13 |
| December | 4 |
